# Variations in the frequency and amplitude of resting-state EEG and fMRI signals in normal adults: The effects of age and sex

**DOI:** 10.1101/2020.10.02.323840

**Authors:** Xiaole Zhong, J. Jean Chen

## Abstract

Frequency and amplitude features of both resting-state electroencephalography (EEG) and functional magnetic resonance imaging (fMRI) are crucial metrics that reveal patterns of brain health in aging. However, the association between these two modalities is still unclear. In this study, we examined the peak frequency and standard deviation of both modalities in a dataset comprising healthy young (35.5±3.4 years, N=134) and healthy old (66.9±4.8 years, N=51) adults. Both age and sex effects were examined using non-parametric analyses of variance (ANOVA) and Tukey’s Honest Significant Difference (HSD) post-hoc comparisons in the cortical and subcortical regions. We found that, with age, EEG power decreases in the low frequency band (1-12 Hz) but increases in the high frequency band (12-30 Hz). Moreover, EEG frequency generally shifts up with aging. For fMRI, fluctuation amplitude is lower but fluctuation frequency is higher in older adults, but in a manner that depends on the fMRI frequency range. Furthermore, there are significant sex effects in EEG power (female > male), but the sex effect is negligible for EEG frequency as well as fMRI power and frequency. We also found that the fMRI-EEG power ratio is higher in young adults than old adults. However, the mediation analysis shows the association between EEG and fMRI parameters in aging is negligible. This is the first study that examines both power and frequency of both resting EEG and fMRI signals in the same cohort. In conclusion, both fMRI and EEG signals reflect age-related and sex-related brain differences, but they likely associate with different origins.

## Introduction

Understanding the effects of healthy aging on brain oscillations may help explain many of the observed cognitive age effects. Techniques such as electroencephalography (EEG) and magnetoencephalography (MEG) can provide measurements more closely reflecting the underlying neuronal signals. In the alpha EEG, which is the most widely studied EEG band in aging, the effect of age on oscillatory power has been widely studied (see review in (Ishii et al., 2017)). The age effect on the other EEG bands, including the slow-wave delta and theta bands, and the fast-wave beta band, is less well studied. Decline in slow-wave EEG power in aging has been associated with deficits in perceptual speed and executive function (Vlahou et al., 2014), while enhancement in fast-wave beta EEG power in aging is associated with reduced specificity in motor processing and imagery (Christov and Dushanova, 2016a). EEG oscillatory frequency can also be affected by processes in aging. EEG and MEG studies have reported a reduction of neuronal activity frequency in mild cognitive impairment (Garcés et al., 2013). Hence, the age-associated slowing of brain activity may provide an early indication of impending dementia. However, frequency measurements have thus far been limited to EEG and MEG studies.

Given the limited spatial resolution of EEG and MEG, the use of resting-state functional MRI (rs-fMRI) for brain-oscillation mapping is increasingly common. Metrics such as brain variability (Garrett et al., 2010; Kumral et al., 2019), the amplitude of low-frequency fluctuations (ALFF) (Zou et al., 2008) and the resting-state fluctuation amplitude (RSFA) (Kannurpatti et al., 2012) are all indicators of oscillatory amplitude. The rs-fMRI approach offers more comprehensive spatial information that can be easily integrated with structural MRI data. These variables are associated with vascular (Tsvetanov et al., 2015), metabolic (Jiao et al., 2019) and cognitive influences (Garrett et al., 2017). However, the findings have been varied, and the interpretation of these metrics remains ambiguous.

Sex differences in rs-fMRI functional connectivity have been reported in multiple studies (Hjelmervik et al., 2014; Zhang et al., 2016), generally with males showing higher connectivity than females, but sex effects on the underlying fMRI signal fluctuation amplitude and EEG oscillation power are scarcely reported. Still more scarce is reporting on the interaction between age and sex in shaping the EEG and fMRI metrics.

The above mentioned research beckon the question “what is the relationship between alterations in electrophysiological and rs-fMRI oscillations in aging?”. Using data from the Leipzig mind-brain-body (LEMON) study, Kumral et al. recently published on age and sex differences in EEG and fMRI fluctuation amplitudes in healthy adults (Kumral et al., 2019). The study produced the first direct comparison of resting-state effects of age and sex on EEG and fMRI power metrics, acquired from the same subjects, and concluded that age effects on fMRI amplitude were not related to those found in EEG power. Nevertheless, some questions remain unanswered. First, shifts in EEG frequency in aging have been reported, and it is unclear whether frequency shifts occur in rs-fMRI in aging. Second, it is unclear if there is a difference between the sexes in how EEG and rs-fMRI frequencies vary with age. Third, it is unclear whether age-related rs-fMRI frequency variations are mediated by those in EEG.

In this study, we address these gaps in knowledge by extending analysis of data from the LEMON study. We hypothesize that: (1) the peak frequency of all EEG bands differ significantly across age and sex groups; (2) the peak frequency of the rs-fMRI signal is lower in the older adults, but there should be no difference between sexes; (3) as the neuronal contribution to the fMRI signal varies with fMRI frequency, the age and sex effects of the rs-fMRI signal amplitude and frequency vary depending on the fMRI frequency band in question.

## Methods

### Participants

The “Leipzig Study for Mind-Body-Emotion Interaction” (LEMON, publicly available at: http://fcon_1000.projects.nitrc.org/indi/retro/MPI_LEMON.html) (Babayan et al., 2019) dataset comprises 227 healthy subjects in two age groups. The older group is aged between 59-77 years old (N=74, 37 females) while the younger group between 20-35 years old (N=153, 45 females). No participant reported a history of cardiovascular disease, psychiatric disease, neurological disorders, malignant disease, or medication/drug use that could affect the study. The study protocol conformed to the declaration of Helsinki and was approved by the ethics committee at the medical faculty of the University of Leipzig (reference number 154/13-ff).

We performed additional QC and excluded data sets that had incomplete data, mismatching sampling rates, image artefacts, excessive head movement, or excessive background noise. The final sample with 185 subjects used in this study included 134 young (20-35 years old, 42 females) and 51 old subjects (59-77 years old, 23 females). Ages were only recorded by the LEMON study quinquennially (5 year steps), thus the group means and standard-deviations of the mean are provided. For age distributions see Figure A1 in the Supplementary Materials. Furthermore, the education levels were provided in the German system (Hauptschule and above), and were not translatable into years of education.

### Data acquisition

All data acquisitions are described in the LEMON paper ((Babayan et al., 2019). The relevant sections are provided below.

#### EEG

Sixteen minutes of resting-state EEG was recorded with BrainAmp MR-plus amplifiers using 62-channel (61 scale electrodes and 1 VEOG electrode below the right eye) active ActiCAP electrodes (both Brain Products GmbH, Gilching, Germany) attached according to the international standard 10-10 system and referenced to FCz. The ground electrode located at the sternum and skin-electrode interface impedance was kept below 5 kOhm. The EEG signal is digitized at a sampling frequency 2500 Hz and amplitude resolution was set to 0.1 micro-Volts. The EEG session included a total of 8 eyes-closed (EC) blocks and 8 eyes-open (EO) blocks, each 60 s. Subjects were asked to fixate on a black cross on a white background during the EO period, demonstrated using Presentation software (Version 16.5, Neurobehavioral System Inc., Berkley, CA, USA). As rsfMRI data were collected only in the EO condition, only EEG from EO the condition was used in the comparative analysis.

#### MRI

On a separate day from the EEG session, magnetic resonance imaging (MRI) was performed on a 3 Tesla scanner (MAGNETOM Verio, Siemens Healthcare GmbH, Erlangen, Germany) equipped with a 32-channel head coil. Participants were informed to keep their eyes open while focusing on a low-contrast cross during the scan.

Structural T1-weighted image was acquired using an MP2RAGE sequence with parameters: TR=5000 ms, TE=2.921 ms, TI1=700ms, TI2 =2500ms, FA1=4 deg, FA2=5 deg, bandwidth=240 Hz/pixel, FOV=256 mm, voxel size 1 mm isotropic, 176 slice encodes. Functional image recorded with T2*-weighted gradient-echo EPI sequence parameters with parameters, TR=1400 ms, TE=30 ms, flip angle=69 deg, bandwidth=1776 Hz/pixel, partial Fourier ⅞, voxel size 2.3 mm isotropic, 64 slices.

### EEG data preprocessing and analysis

The EEG processing strategy is summarized in Fig. 1a. EEG preprocessing was conducted with EEGLAB (version 14.1.1bl Delorme and Makeig, 2004) functions implemented in Matlab (The MathWorks Inc. Natick, Massachusetts, USA). The raw EEG data were down-sampled from 2500 Hz to 250 Hz, band-pass filtered within 1-45 Hz with 4^th^ order back and forth Butterworth filter before the split into EO and EC conditions. 6.6% of the data were rejected by visual inspection, due to facial muscular tension and gross movements, artifactual channels. Furthermore, principal component analysis was used to reduce the dimensionality of the data by including at least 30 principle components that can explain 95% of the total variance. Then, using independent component analysis (Bell and Sejnowski, 1997), signal components related to physiological sources e.g. eye blinks, eye movements, residual ballistocardiograph artefacts and muscle activity were rejected. Preprocessed EEG signals were re-referenced to the common average and channel FCz was added as a normal channel.

**Figure 1.**
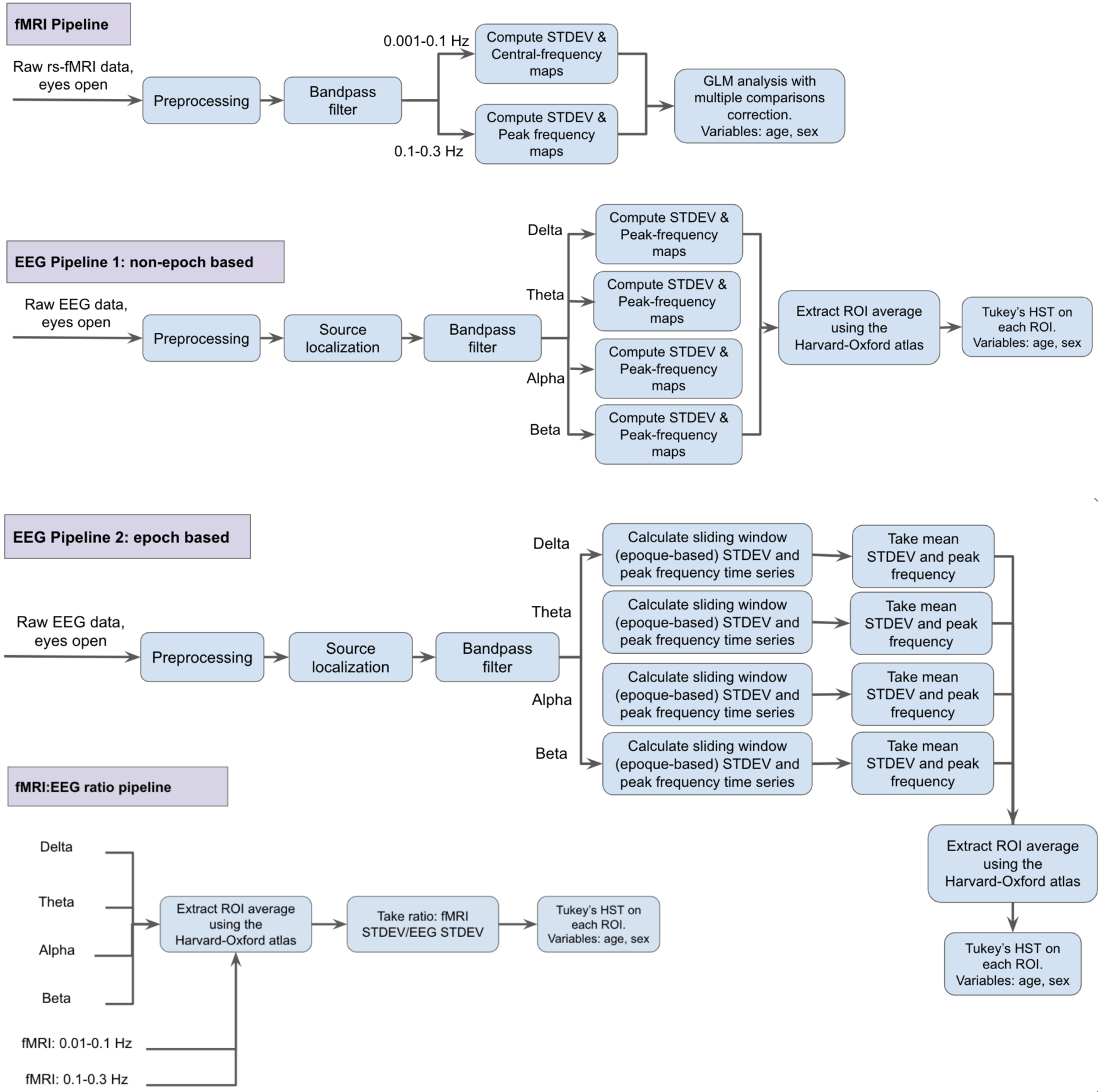
Overview of analysis procedure. Two EEG pipelines and one fMRI pipeline are used.

The geometry of the source reconstruction model was based on the MNI/ICBM152 (International Consortium for Brain Mapping) standard anatomy. eLORETA (exact low-resolution brain electromagnetic tomography) implemented in the M/EEG toolbox of Hamburg (Haufe and Ewald, 2019; Pascual-Marqui, 2007) METH; (Haufe and Ewald, 2019; Pascual-Marqui, 2007) was used to compute source distribution from the scalp EEG recordings. The leadfield matrix was generated to relate 2113 source voxels and 62 scalp electrodes. Singular value decomposition of each voxel was used to determine the dominant orientation of the source signal, followed by bandpass filters to filter EEG signal into specific frequency bands, associated with brain oscillations: delta (1-4 Hz), theta (4-8 Hz), alpha (8-12 Hz), and beta (12-30 Hz).

For the non-epoch based approach (Fig. 1b), the peak frequency in each EEG band was calculated by the center-of-mass approach. This approach ensured that the frequency characteristic was more robust against noise than when identifying a single peak. The amplitude envelope of each band’s oscillations was extracted using the Hilbert transform (Rosenblum et al., 2001) and then temporally smoothed by a kernel of full-width at half-maximum (FWHM) 0.5 s. Power was computed as the standard deviation (STDEV) of the each source’s smoothed time series, which are further averaged within 106 regions of interest (ROIs) as designated according to the Harvard-Oxford subcortical and cortical atlas (Desikan et al., 2006). For the epoch-based approach (Fig. 1c), the STDEV were calculated for moving epochs that match the TR of the fMRI data (1.4 sec). This step attempts to further match the temporal features in the EEG and fMRI time series, and is analogous to the epoch-wise variability measure introduced by Garrett et al. (Garrett et al., 2010). The standard deviation and peak frequency of each epoch was computed by the same method as the non-epoch based approach. Subsequently, the standard deviation of each data set is calculated by the standard deviation all epoch-specific standard deviations. Due to limited data points in epoch based approach, peak frequencies were defined by the average of maximum power frequency in each epoch.

### rs-fMRI data preprocessing and analysis

The rs-fMRI processing strategy is summarized in Fig. 1. fMRI preprocessing was implemented with tools from FSL (Jenkinson et al., 2012) and FreeSurfer (Fischl, 2012). The following steps were included in preprocessing: (I) 3D motion correction (FSL mcflirt), (II) slice-timing correction (FSL slicetimer), (III) brain extraction (FSL bet2 and FreeSurfer mri_watershed), (IV) Rigid body coregistration of functional data to the individual T1 image (FSL flirt), (V) Regress out the effect of artifacts (fsl_glm), (VI) bandpass filtering to obtain band 1 (0.01-0.1 Hz) and band 2 (0.1-0.3 Hz), (VII) spatial normalization to MNI152 (Montreal Neurological Institute) standard space with spatial resolution 2mm isotropic (FSL flirt), (VIII) the data were spatially smoothed with 6 mm full-width half-maximum Gaussian kernel (FSL fslmaths). The two frequency bands are meant to capture fluctuations that are typically associated with neuronal activity (lower frequency) and physiological processes (higher frequency). The bandpass filter was implemented using Matlab (327th order Kaiser bandpass FIR filter with respective passband) to ensure minimal overlap between the bands.

Standard deviation and peak frequency were calculated within each voxel time series then segmented to 106 ROIs using the Harvard-Oxford subcortical and cortical atlas (Desikan et al., 2006) for the mediation and power-ratio analyses.

Furthermore, the ratio of the rs-fMRI and EEG signal fluctuations is taken between signal pairs across the 2 fMRI frequency bands and 4 EEG bands. This measurement is intended to produce a surrogate of the vascular-neuronal fluctuation ratio in the resting state.

## Statistical methods

To investigate age and sex effect, log-transformed EEG standard deviation and peak frequency from each 115 ROIs were examined by non-parametric analyses of covariance (ANCOVAs, type III) after outliers (data points out of 1.5 interquartile range) were removed. The significance of group differences were further tested by Tukey’s Honest Significant Difference (HSD) post-hoc comparisons with a significance threshold 0.05. Moreover, age-biases in the sex differences were assessed by performing the sex-group comparisons for young and older adults separately, and sex biases in age differences were assessed by performing age-group comparisons for men and women separately. All analyses were implemented in Matlab (The MathWorks Inc., Natick, Massachusetts, USA).

BOLD fMRI STDEV and peak frequency were tested for effects of age and sex with the randomized design generalized linear model (GLM) implemented in FSL (Jenkinson et al., 2012) with significance threshold 0.05, corrected for multiple comparisons using randomise. Subcortical region standard deviation and peak frequency were also examined by non-parametric analyses of covariance (ANCOVAs, type III) and Tukey’s HSD post-hoc comparisons to better investigate age and sex effects in these ROIs.

Two statistical approaches were used to reveal the relationship between BOLD fMRI and EEG. The first approach used non-parametric analysis of covariance (ANCOVAs, type III) and Tukey’s HSD post-hoc test to analyze the age and sex effect on fMRI-EEG power ratio. The second is a set of mediation analyses.

Mediation pathway analysis (Hayes, 2013) was applied to investigate the association between EEG frequency and BOLD frequency as well as EEG standard deviation and BOLD standard deviation. The models, summarized in Fig. 2, were built in Matlab (The MathWorks Inc., Natick, Massachusetts, USA) with the Variational Bayesian Analysis (VBA) toolbox (Daunizeau et al., 2014). EEG parameters were set as a mediator while age or sex and BOLD parameters were the independent and dependent variable, respectively. The model was first built through Baron & Kenny’s 3-step mediation analysis (Baron and Kenny, 1986), then examined by the Sobel test (Sobel, 1982) to test if the relationship between the independent variable and dependent variable is significantly reduced when including the mediator. Significance was declared when the p-value from the Sobel test was lower than 0.05. A pairwise t-test which corrected for false discovery rate was added after mediation analysis to examine the Sobel test (Sobel, 1982) p-value of epoch based and non-epoch based mediation analysis results.

**Figure 2.**
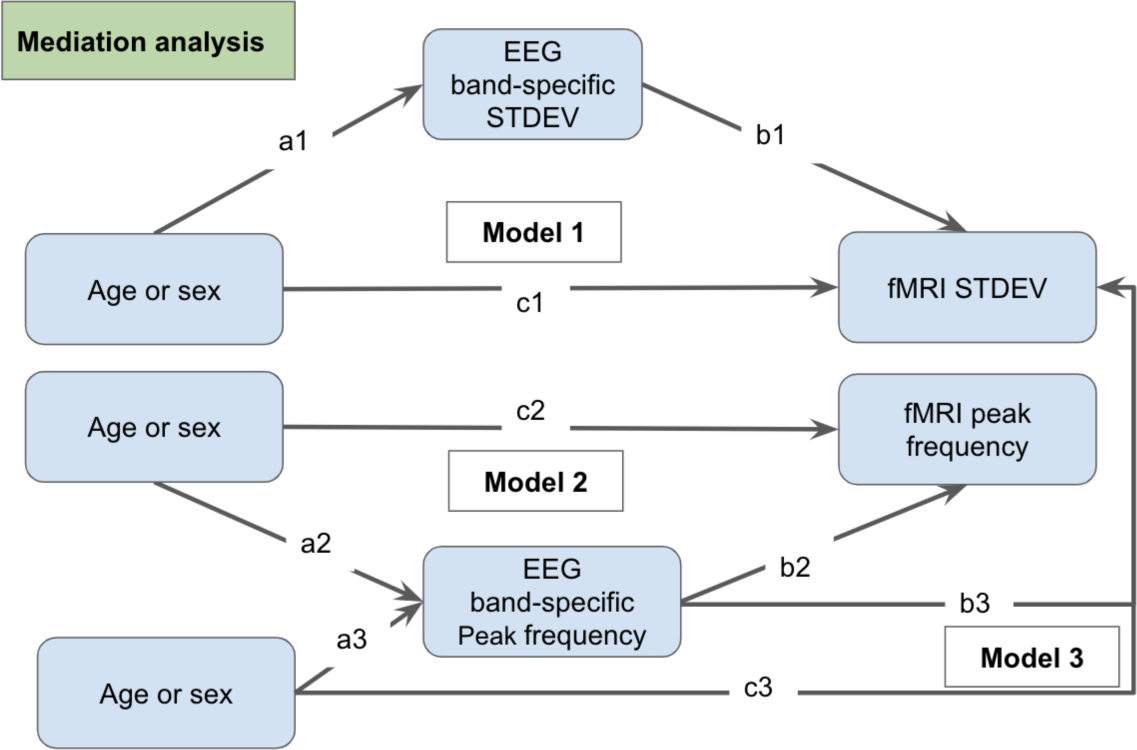
Models for the mediation analyses. We investigate the mediating effects of EEG power (Model 1) and EEG frequency (Models 2 & 3) on rs-fMRI amplitude (STDEV) and frequency. The mediation coefficients a, b, and c represent indirect and direct effects. The independent variables are age and sex.

## Results

### EEG Power

The graphical distribution of the significant cortical and subcortical age effects are shown in Fig. 3 and Fig. 7, respectively. For the non-epoch based approach, Tukey’s HSD post-hoc comparison shows that younger subjects exhibit lower EEG signal power in the delta, theta and alpha band. Significant main effects for the delta band are found in 31 ROIs in parietal, frontal, and limbic regions and 8 ROIs in the subcortical region, while theta band shows significant age differences in 13 ROIs in the limbic and frontal cortex, and no significant effects in subcortical ROIs. In addition, for the alpha band, 9 ROIs in the occipital lobe show significant age effects but no subcortical ROI shows significant effects. The beta band shows a different trend than the other three bands; 28 ROIs in frontal and temporal regions show a significant increase with brain aging.

**Figure 3.**
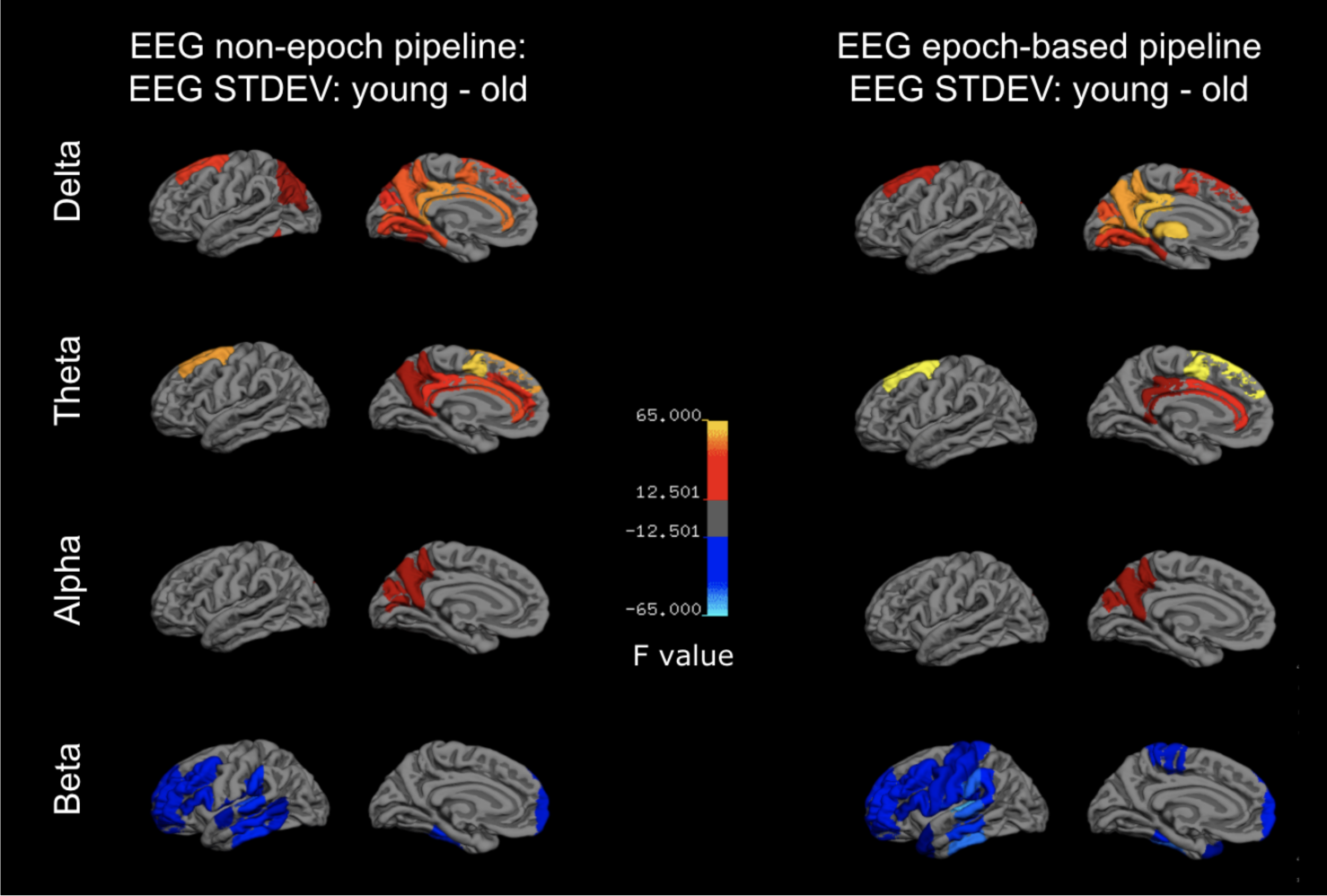
EEG power versus age. All differences correspond to young minus old, and only statistically significant results are shown in colour. EEG power is computed as the signal standard-deviation, using either the epoch-based or non-epoch pipeline. Delta and theta power are negatively associated with age in the superior frontal gyrus, the anterior and posterior cingulate and the precuneus. Alpha power is negatively associated with age only in the posterior cingulate and precuneus, while beta power is positively associated with age, being higher in the older adults in the frontal and temporal lobes. There was no substantial difference between non-epoch and epoch-based pipelines except for in the beta band, where the age-affected region is expanded.

The significant cortical and subcortical sex effects are shown in Fig. 4 and Fig. 8, respectively. Concerning sex effects, females exhibit higher EEG power in the delta band (14 ROIs in occipital and limbic regions), theta band (28 ROIs in frontal, limbic, and occpital cortices), the alpha band (5 ROIs in the frontal lobe), and beta band (76 ROIs across the brain).

**Figure 4.**
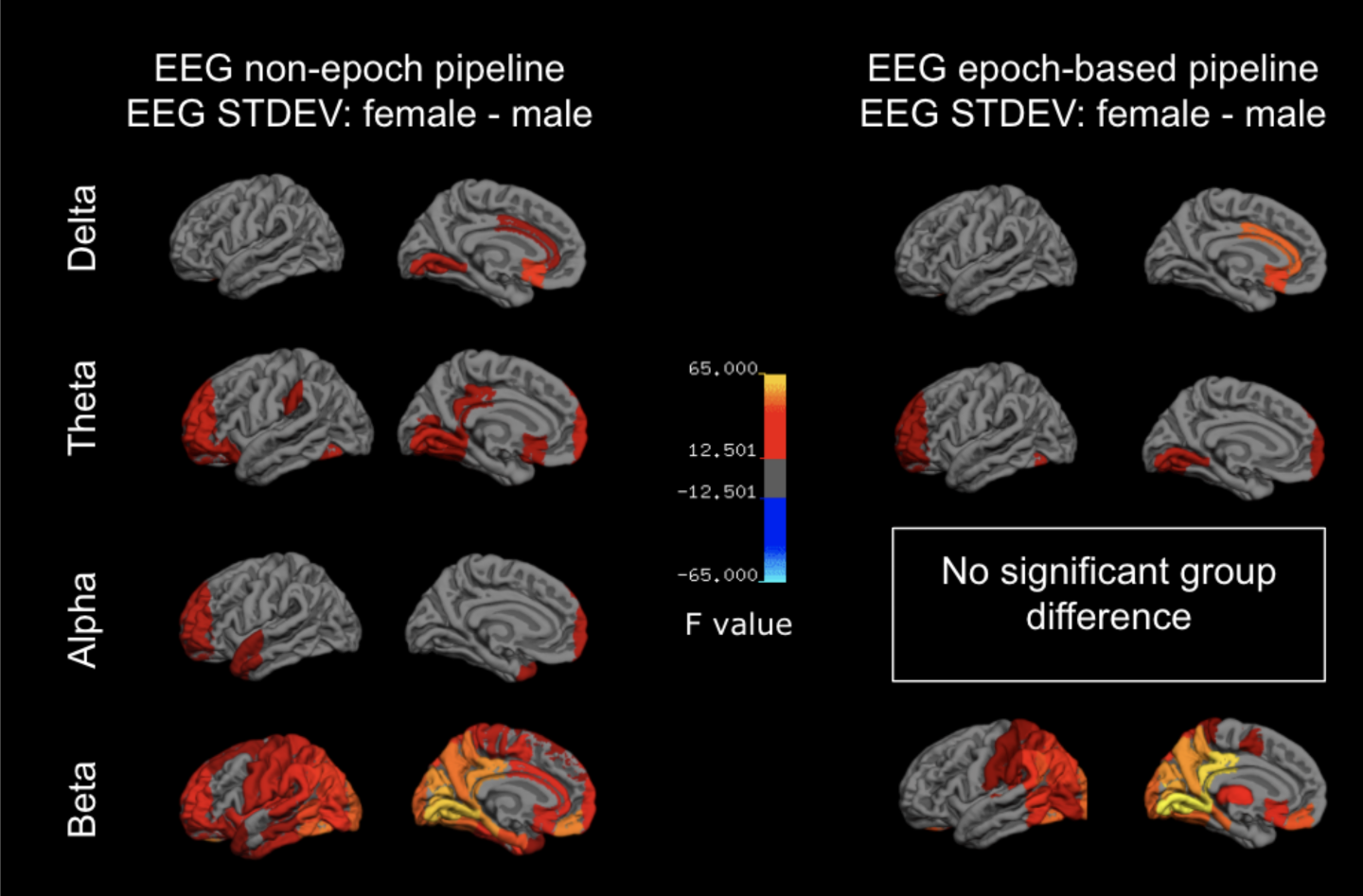
EEG power versus sex. All differences correspond to females minus males, and only statistically significant results are shown in colour. EEG power is computed as the signal standard-deviation, using either the epoch-based or non-epoch pipeline. In the delta band, women have higher EEG power than men in the superior frontal and cingulate cortices. In both the alpha and theta bands, EEG power is higher in women in the frontal and temporal regions. In the beta band, the differences are the most widespread, covering almost the entire cortex (with the exception of the temporal regions showing higher alpha power in women). In the epoch-based pipeline, the effects are less pronounced. Notably, the delta band still shows higher EEG power in women in the cingulate cortex, as in the non-epoch case, but in the beta band, this effect is limited to the posterior portion of the cortex.

**Figure 5.**
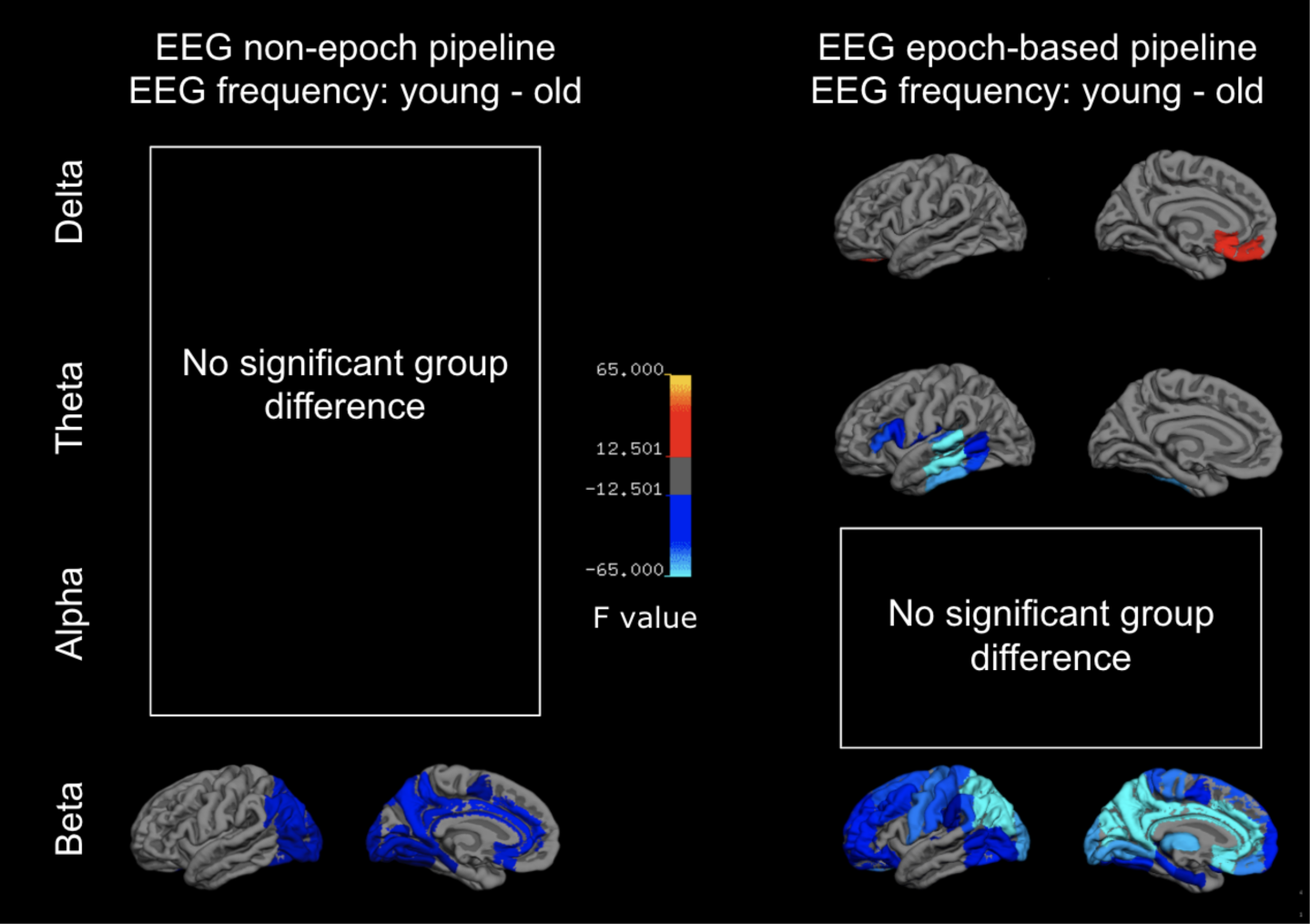
EEG frequency versus age. All differences correspond to young minus old, and only statistically significant results are shown in colour. In the non-epoch approach, beta band is the only band whose frequency is significantly associated with age, being higher in older adults in the cingulate, precuneus and occipital lobe. The epoch-based approach shows more age differences in frequency, with the delta band showing higher frequency in young adults and theta band showing lower frequency in young adults. In the beta band, the frequency difference is higher in older adults in nearly the entire cortex (with the exception of the temporal regions showing higher theta frequency in older adults).

**Figure 6.**
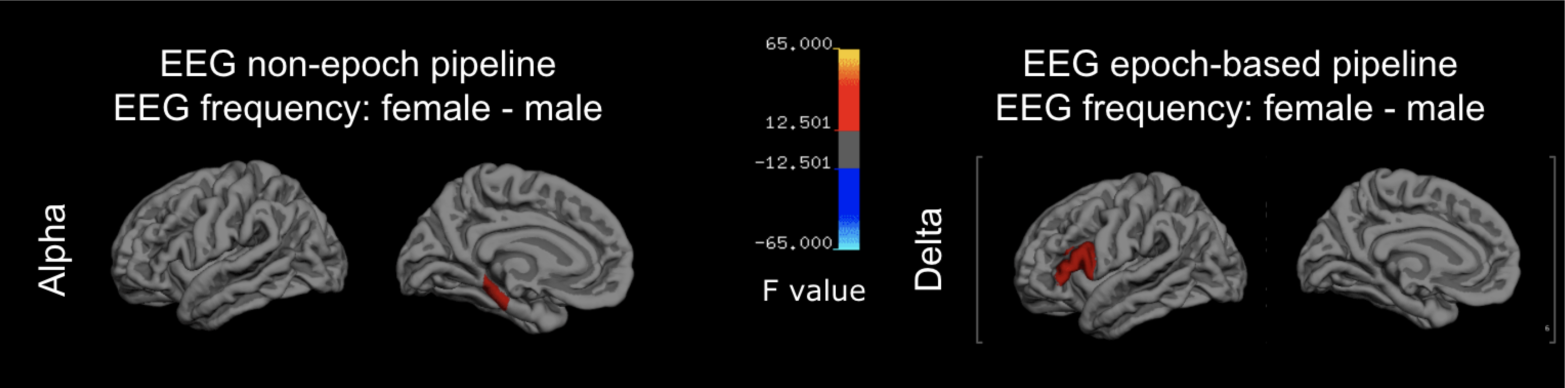
EEG frequency versus sex. All differences correspond to young minus old, and only statistically significant results are shown in colour. There is minimal difference in EEG frequency between men and women (in the frontal gyrus), using either EEG processing pipeline.

Epoch-based approach analysis shows less significant ROIs for age and sex effects on EEG power than the non-epoch based EEG time series. As with the non-epoch based results, younger participants have higher EEG signal power than older participants in the alpha band (7 ROIs in the occipital lobe), theta band (8 ROIs in limbic and frontal regions), and delta band (19 ROIs in parietal, frontal, and limbic lobes) in the cortical region. Meanwhile, older subjects have higher beta power in frontal and temporal lobes. There is no ROI that shows significant age effect for the alpha band, delta band, and theta band in the subcortical region, while delta-band power shows a significant increase in 2 ROIs. Regarding sex differences for epoch-based EEG power, females have higher standard deviation than males in delta band(13 ROIs), theta band (17 ROIs), and beta band (47 ROIs).

### EEG Frequency

The graphical distribution of the significant cortical and subcortical age effects are shown in Fig. 5 and Fig. 8, respectively. The beta band is the only band in the non-epoch based signal that exhibits a significant age effect for the peak frequency, with 40 cortical ROIs and 10 subcortical ROIs showing a significant increase in peak frequency from young to old. In terms of the sex effect, females have a higher peak frequency than males in the right parahippocampal gyrus posterior division.

**Figure 7.**
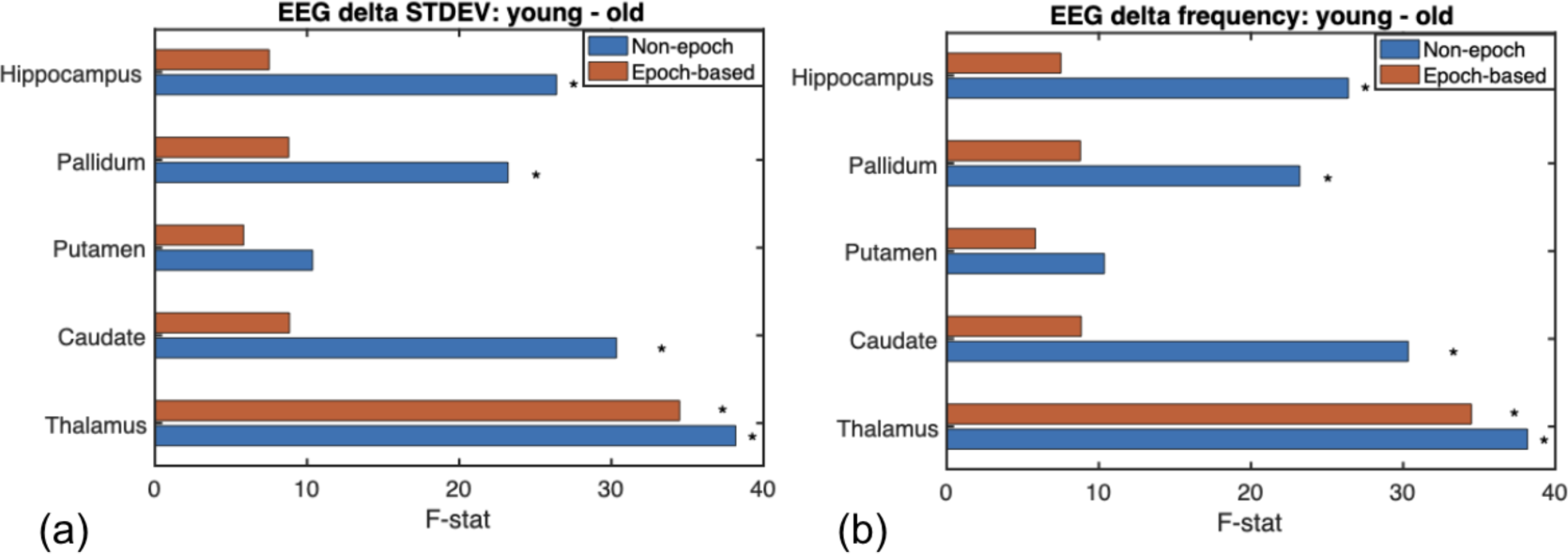
EEG power and frequency versus age: subcortical regions. (a) Only the delta band shows a significant age effect in power (young > old), namely in the thalamus, caudate, pallidum and hippocampus. (b) Delta band is also the only one to show significant age differences in EEG frequency, found in the same four regions. Non-epoch-based age-differences are greater than epoch-based ones. Significance is indicated by asterisks.

**Figure 8.**
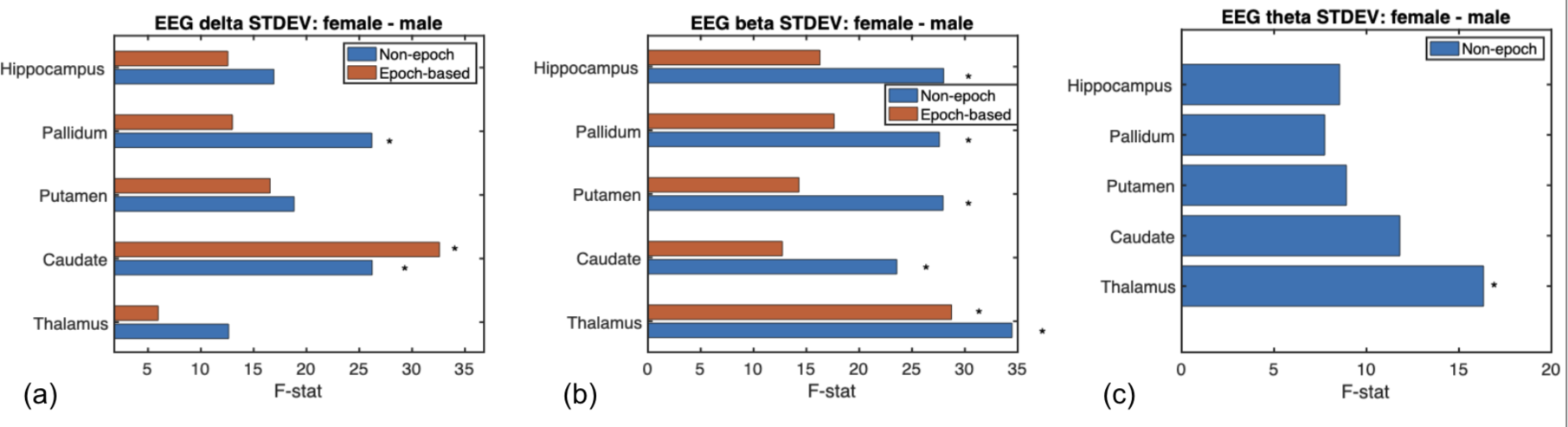
EEG power and frequency versus sex: subcortical regions. (a) Delta, theta and beta bands all show significant sex differences (female > male). (a) For delta band, the difference is only in the caudate and pallidum, whereas (b) for beta band, it extends to the thalamus, putamen and hippocampus as well. For the theta band, the difference is only in the thalamus. In all cases, non-epoch-based age-differences are greater than epoch-based ones. Significance is indicated by asterisks.

The epoch-based approach shows significant age effect in the delta band, theta band, and the beta band in the cortex. Delta band peak frequency shows a significant reduction in the older adults in 6 ROIs, while theta and beta band show a significant increase with brain aging in 20 ROIs and 21 ROIs, respectively. Epoch-based EEG peak frequency only shows a significant age effect in the cortical region. There is no significant frequency age effect in any subcortical ROI for the delta and theta bands, but beta band frequency is significantly lower in older adults in 10 subcortical ROIs. Delta band is the only band that shows a significant sex effect for epoch-based EEG peak frequency -- females have higher peak frequency than males in the left and right frontal gyrus.

### fMRI Fluctuation Amplitude

From Fig. 9, it is evident that young adults have qualitatively higher fMRI signal amplitudes and lower fMRI frequencies than older adults. Moreover, older adult amplitude (STDEV) maps display lower inter-regional variability than those of younger adults. rs-fMRI signal fluctuation amplitude is also significantly lower in older adults. For the lower frequency BOLD signal (0.01 to 0.1 Hz), younger adults show higher STDEV in the cingulate, superior frontal gyrus, middle frontal gyrus, and lingual gyrus for the cortical region (Fig. 10). Compared with the 0.01-0.1 Hz BOLD signal, BOLD signal with frequency 0.1 - 0.3 Hz shows more ROIs with significant age effects, which covers most of the cortical regions. For the subcortical region, both frequency ranges show a significant age effect on putamen (Fig. 12). Concerning sex effect, only lower frequency BOLD showed females had significantly higher amplitude and a significant region located in the frontal lobe (Fig. 11).

**Figure 9.**
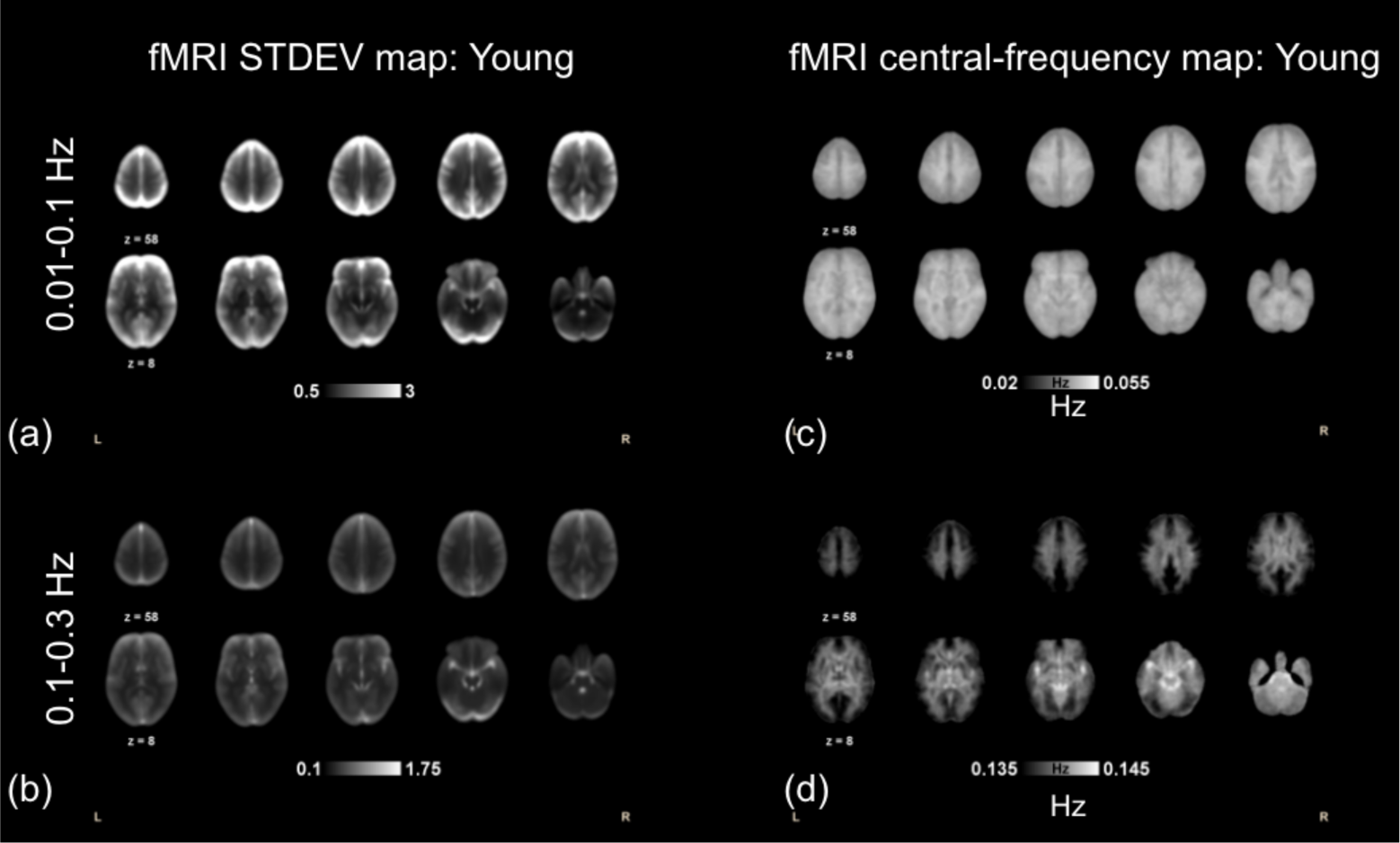
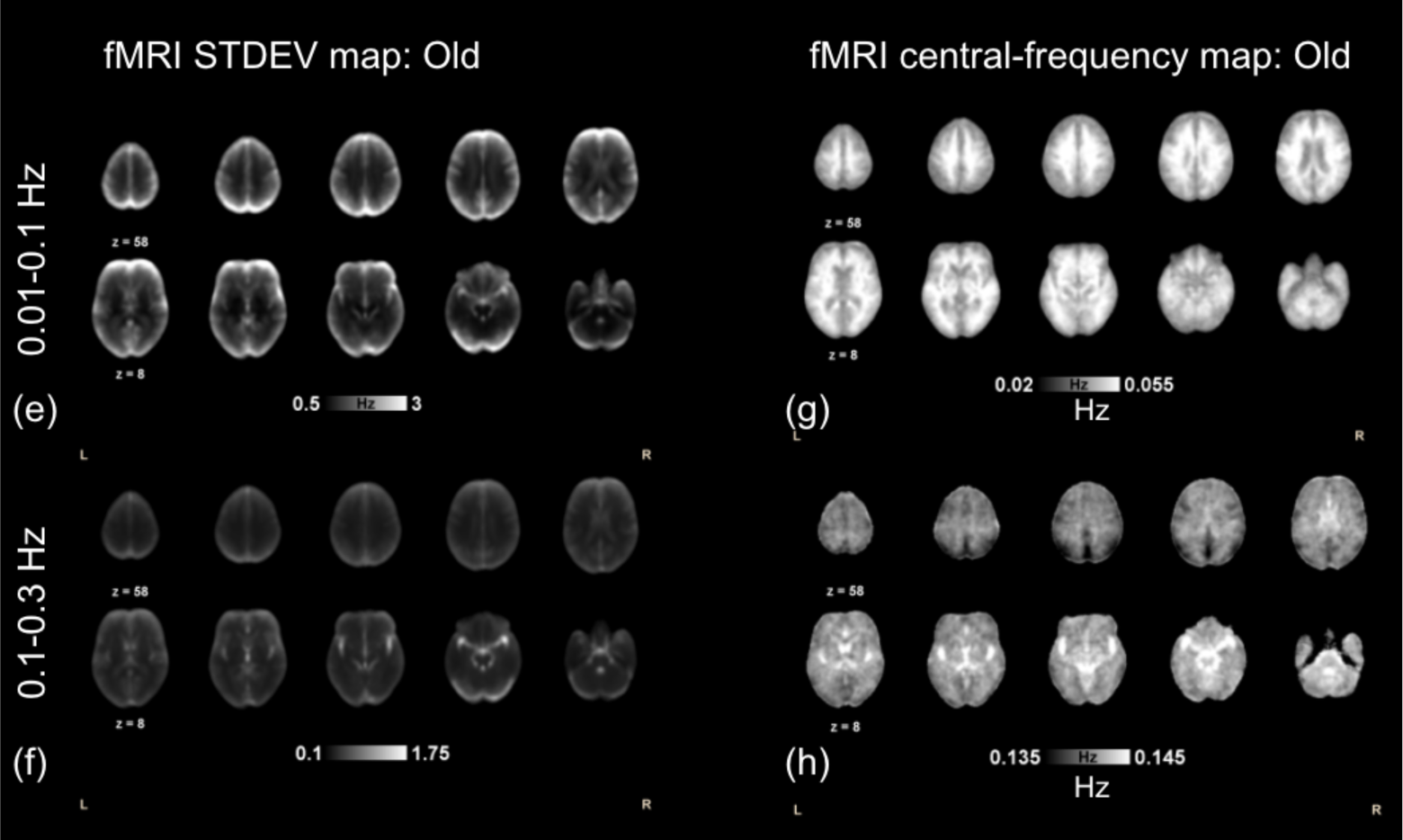
Mean rs-fMRI amplitude and frequency maps in young and old groups. The amplitude is given in units of %ΔBOLD. Cortical regions show higher amplitude (STDEV) than subcortical regions, and young adults (a-d) have higher amplitude but lower frequency than older adults (e-h).

**Figure 10.**
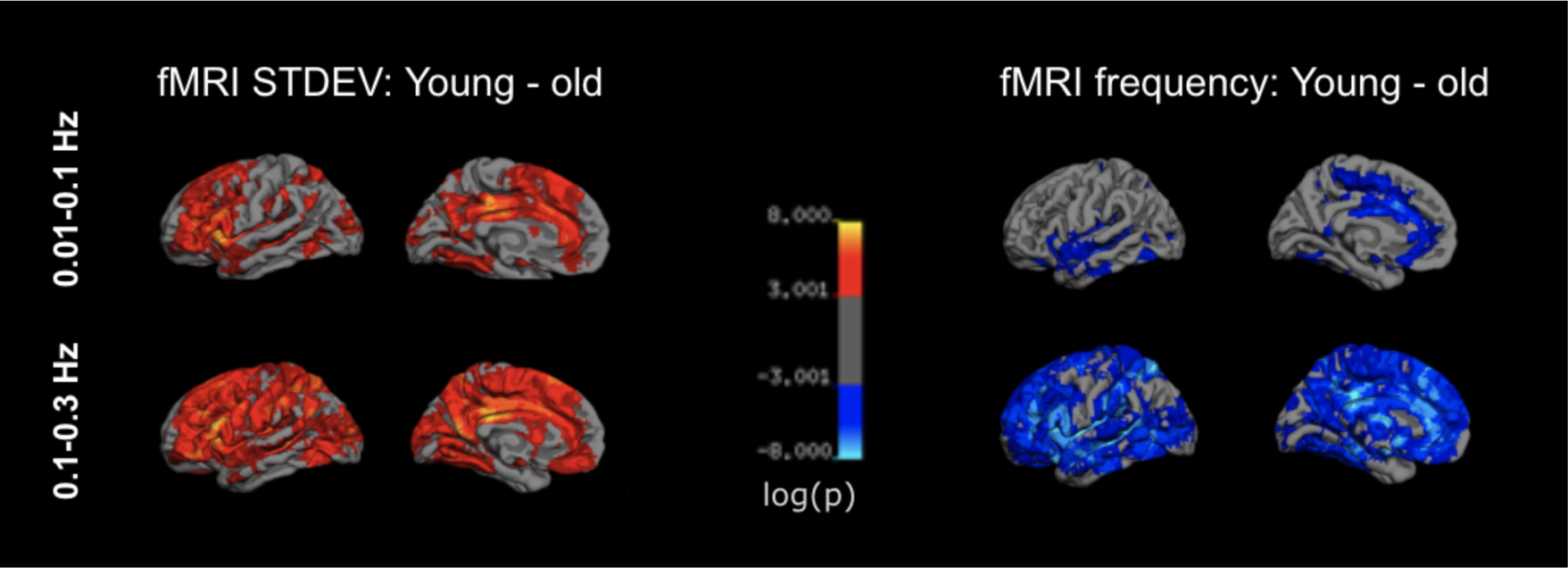
rs-fMRI amplitude and frequency versus age. Only significant regions are shown in colour. fMRI fluctuation amplitude (STDEV) is higher in the younger adults in both the 0.01-0.1 Hz and the 0.1-0.3 Hz bands, with the 0.1-0.3 Hz band showing more widespread differences. fMRI fluctuation frequency is higher in the older adults, also with the 0.1-0.3 Hz band showing more widespread differences. All cases show significant differences in the cingulate cortex and paracentral cortex.

**Figure 11.**
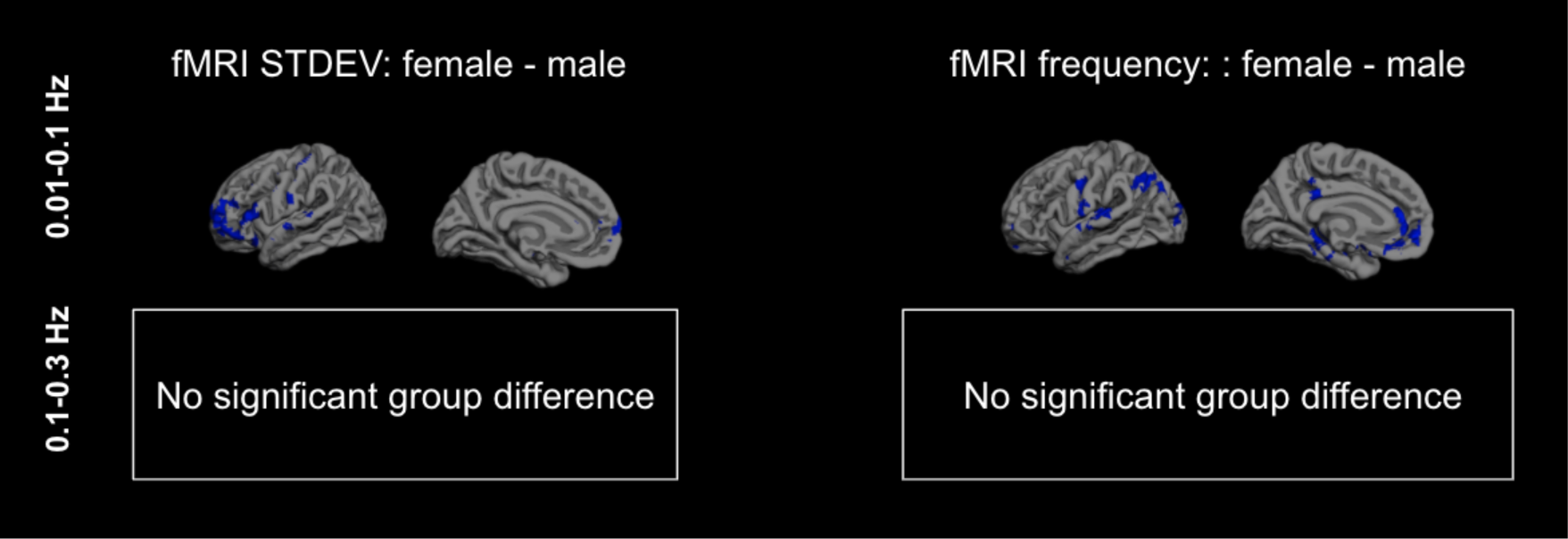
rs-fMRI amplitude and frequency versus sex. All differences correspond to young minus old, and only statistically significant results are shown in colour. There is minimal difference in rs-fMRI frequency between men and women.

**Figure 12.**
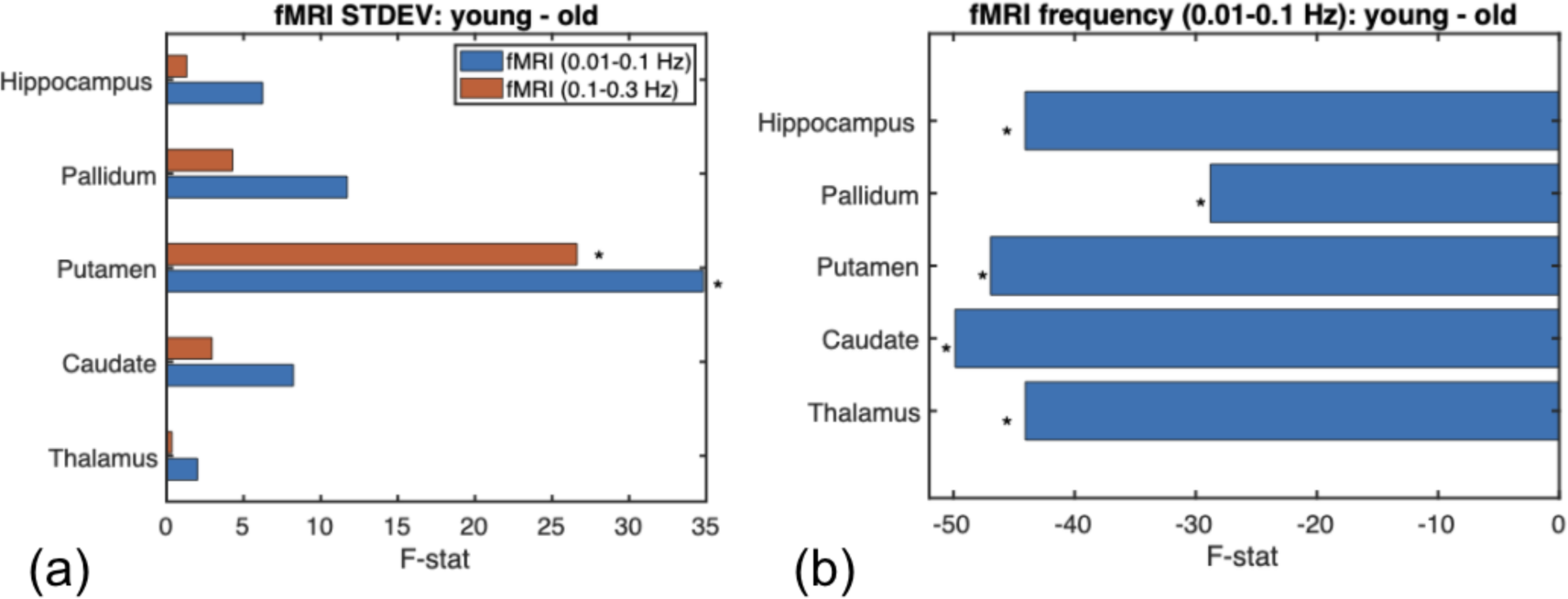
rs-fMRI amplitude and frequency versus age: subcortical regions. (a) The fMRI signal amplitude shows significant age effects (young > old) in the putamen, in both fMRI frequency bands. (b) The fMRI frequency is higher in the older adults in all subcortical structures, but only in the 0.01-Hz band. Significance is indicated by asterisks.

### fMRI Fluctuation Frequency

The peak frequency for 0.01 - 0.1 Hz BOLD signal significantly increases in the superior frontal gyrus, insula, and superior temporal cortex for the cortical region, while the peak frequency for 0.1 Hz to 0.3Hz BOLD signal significantly increases in most of the cortical region (Fig. 10). Regarding the subcortical region, the higher frequency component of the BOLD signal shows older participants have higher peak frequency in the thalamus, caudate, putamen, pallidum, and hippocampus than younger participants (Fig. 12). Males also have higher peak frequency in the occipital and parietal lobe in the cortical region and caudate in the subcortical region for the lower frequency component of the BOLD signal (Fig. 11).

### Age-sex Interactions

As shown in Fig. 14, in terms of the EEG age effects, men exhibit greater age effects in the beta band, while women exhibit greater age effects in the delta and theta bands. Age wise, in the beta and delta bands, older adults exhibit greater sex differences than young adults. The sex biases in the age effects are greater than the age biases in the sex effects. These findings are similar for non-epoch and epoch-based approaches. We did not assess age-sex interactions on the fMRI data, as the effect of sex on fMRI metrics was found to be negligible.

**Figure 13.**
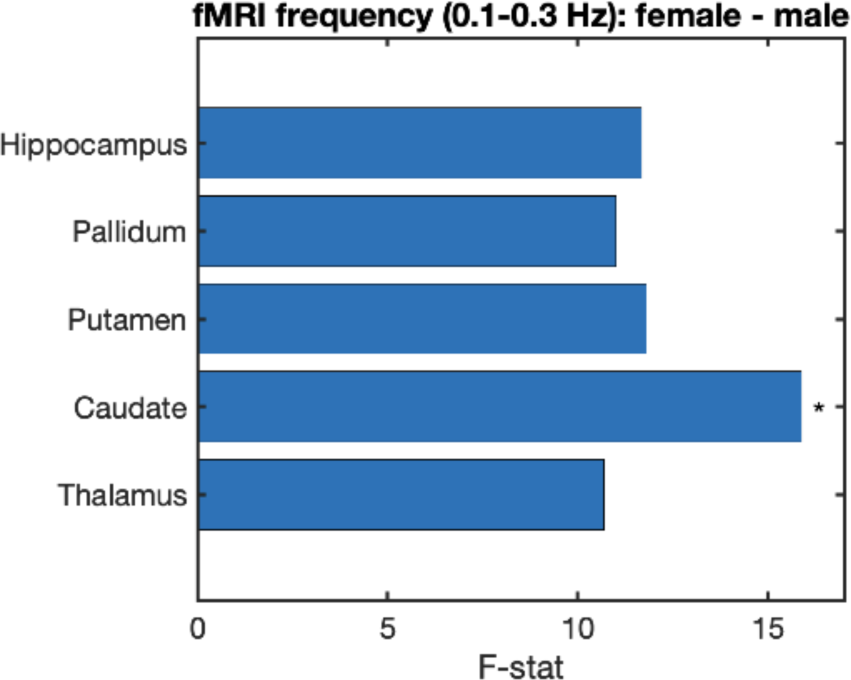
rs-fMRI amplitude and frequency versus sex: subcortical regions. fMRI fluctuation frequency is higher in women only in the caudate, and only in the 0.1-0.3 Hz band. Significance is indicated by asterisks.

**Figure 14.**
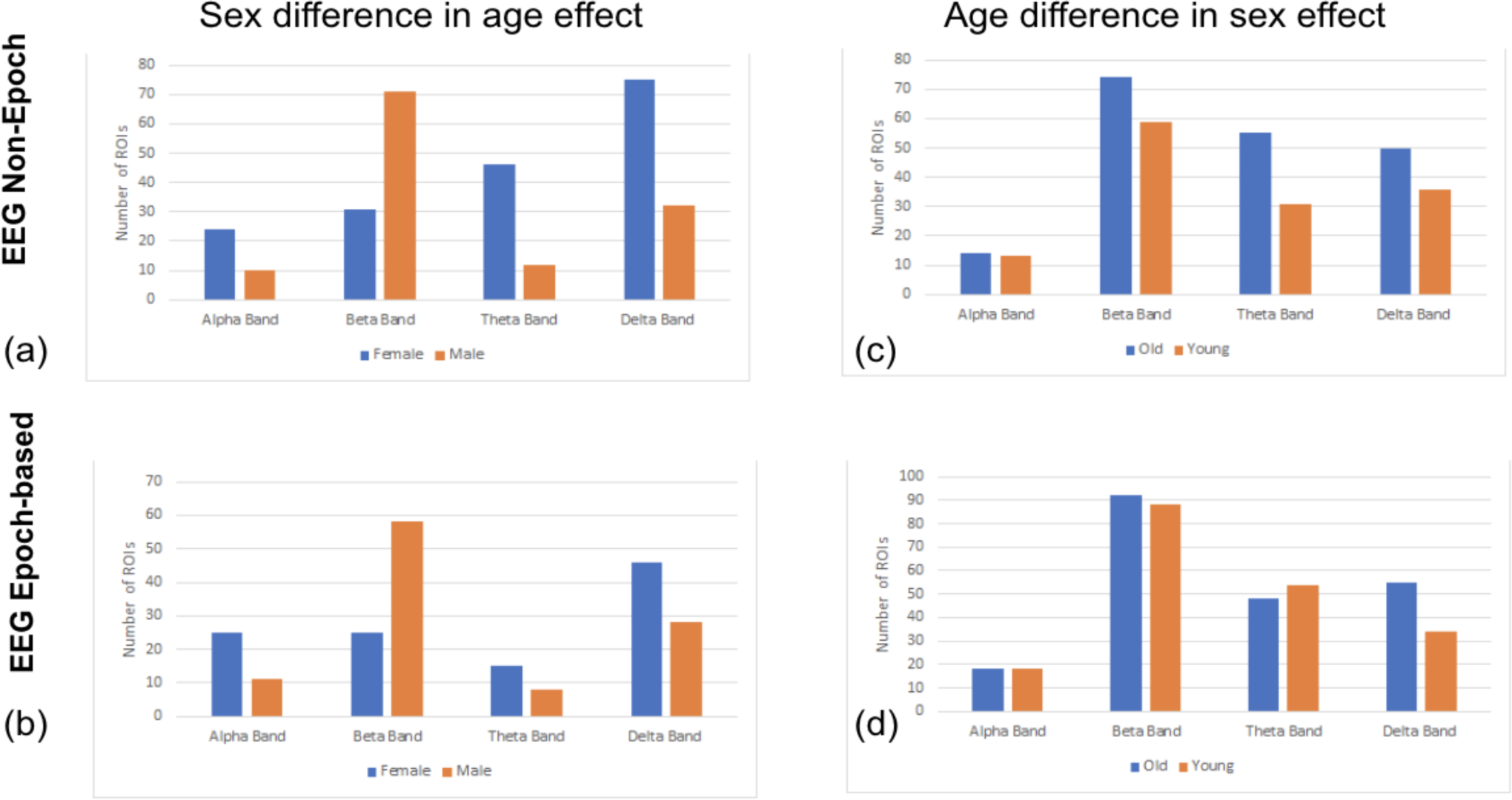
Age and sex interactions in EEG results. Plotted are the numbers of ROIs showing significant age and sex effects. (a) In the beta band, men exhibit more age-group differences (higher orange bar) than women, while women exhibit more age differences in the theta and delta bands. (c) In the beta and delta bands, older adults exhibit greater sex differences than young adults. Both epoch-based (a, b) and non-epoch pipelines (c, d) show similar effects.

### fMRI-EEG Ratios

#### Non-epoch based EEG

The distribution of ROIs showing significant age effects refers to Fig. A2 in Supplementary Materials. The standard deviation ratio between non-epoch based EEG and lower frequency components of the BOLD signal is significantly higher in younger subjects than older subjects in alpha (1 ROI, right inferior frontal gyrus) and beta band (68 ROIs in frontal, parietal, and limbic cortices). In contrast, the ratio is higher for older than for younger participants in the delta band (1 ROI in the thalamus). The ratio between the beta band and higher frequency component of the BOLD signal is the only ratio within 0.1-0.3 Hz BOLD signal is significantly higher in older adults in 71 ROIs in frontal, parietal, and limbic lobes.

The distribution of ROIs showing significant sex effects refers to Fig. A3 in Supplementary Materials. Concerning the sex effect, males shows significantly higher BOLD/EEG standard deviation ratio in all frequency band’s combinations than females. Also, there are more ROIs showing significant sex effects for the fMRI-EEG power ratio in the 0.01-0.1 Hz BOLD band than for in the 0.1-0.3 Hz BOLD frequency band.

#### Epoch based EEG

The distribution of ROIs showing significant age effects is shown in Fig. 15. Statistical analysis reveals that the fMRI-alpha power ratio in the 0.01-0.1 Hz fMRI frequency band is associated with significant age effects (4 ROIs in frontal and parietal regions), being higher in younger adults. The same is true for the fMRI-beta ratio in both the 0.01-0.1 Hz band (61 ROIs in frontal, parietal, and limbic regions) and 0.1-0.3 Hz bands (73 ROIs in frontal, parietal, and limbic regions). The age effect is negligible for the fMRI-delta power ratio in the 0.01-0.1Hz band (1 ROI, medial frontal cortex), the fMRI-alpha band in the 0.1-0.3 Hz frequency band (5 ROIs in frontal and parietal regions) and the fMRI-theta power ratio (1 ROI, middle temporal gyrus).

**Figure 15.**
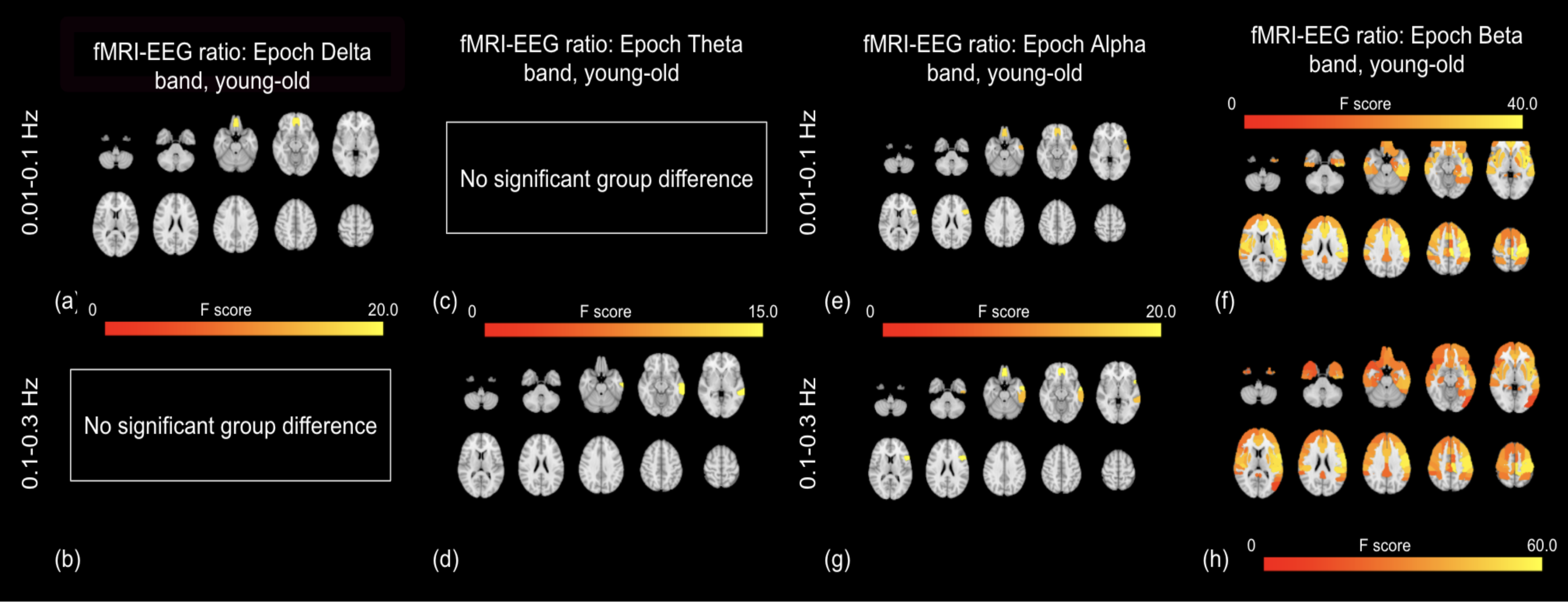
rs-fMRI-EEG power ratio versus age: epoch-based EEG. The fMRI-to-beta power ratio (STDEV ratio) exhibits the greatest age effects (young > old) (f, h). The effects are most pronounced in the 0.1-0.3 Hz range, and the affected regions span nearly the entire anterior cortex, also including subcortical regions such as the putamen and pallidum. The fMRI-to-theta ratio is lower in the young subjects in only the paracentral cortex.

The distribution of ROIs showing significant age effects refers to Fig. 16. Similar to the ratio between BOLD amplitude and non-epoch based EEG power, with females showing significantly lower BOLD-EEG power ratios than males in all frequency combinations.

**Figure 16.**
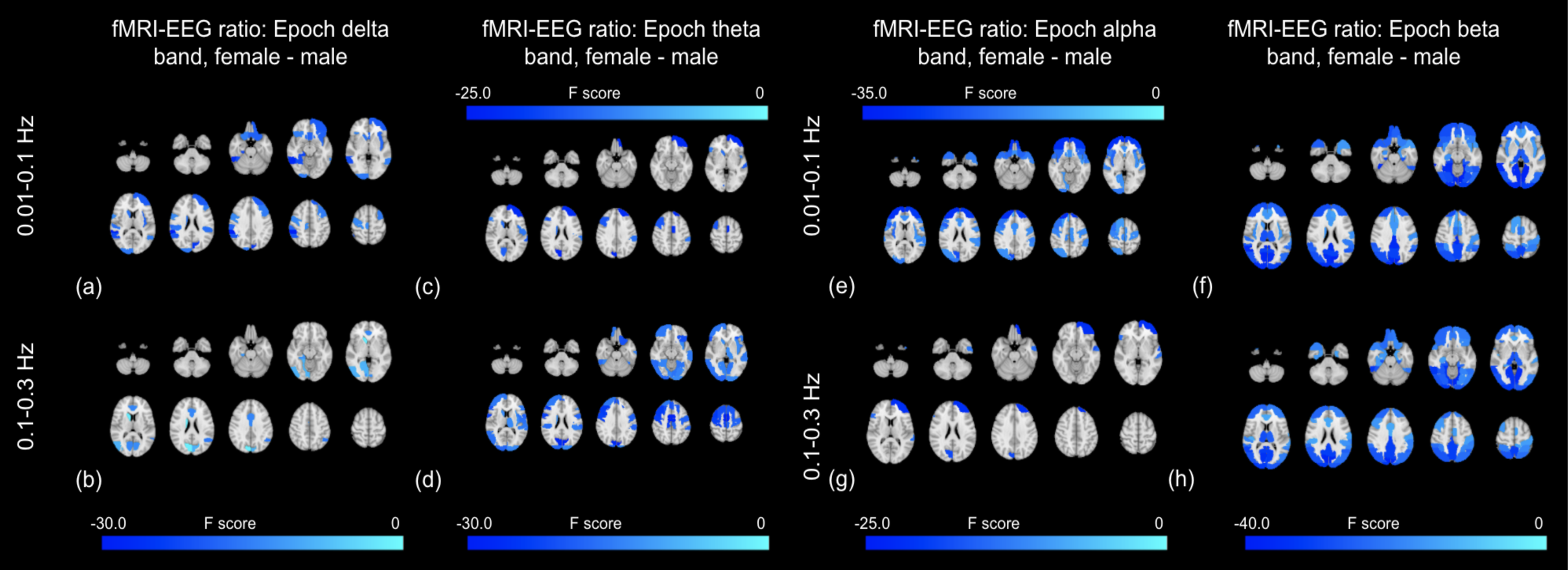
rs-fMRI-EEG power ratio versus sex: epoch-based EEG. Significant sex effects (female < male) are found in the ratios corresponding to all EEG bands, spanning the frontal, occipital and paracentral regions. The effects are greatest for the beta band, and more widespread for the 0.1-0.3 Hz fMRI-to-beta ratio.

### Mediation Analyses

Direct significant age effect is found in 85 ROIs for 0.01-0.1 Hz BOLD STDEV with the average 12.8817% effect and 77 ROIs for Hz BOLD STDEV with the average 11.2038% effect. Direct significant sex effect is found in 8 ROIs for 0.01-0.1 Hz BOLD STDEV with the average 8.2858% effect and there is no significant sex effect for 0.1-0.3Hz BOLD STDEV. Regarding BOLD peak frequency, direct significant age effect is found in 51 ROIs for 0.01-0.1 Hz BOLD peak frequency with the average 8.9799% effect and 95 ROIs for Hz BOLD peak frequency with the average 18.7463% effect. Direct significant sex effect is found in 24 ROIs for 0.01-0.1 Hz BOLD peak frequency with the average 8.2836% effect and there is no significant sex effect for 0.1-0.3Hz BOLD peak frequency.

#### Model 1: EEG STDEV mediating the age or sex differences in fMRI STDEV

The detailed information of ROIs with significant effect refers to Table A1 in Supplementary Materials. The mediation analysis shows non-epoch delta band SD partial mediates 7 ROIs for 0.1-0.3Hz BOLD SD, all in the cortical region. Epoch-based beta band SD partial mediate 2 ROIs in the cortical region and 1 ROIs in subcortical for 0.01-0.1Hz BOLD SD. However, the percentages of mediation are < 1% in all ROIs, so the mediation effects are negligible even for the ROIs indicated as partial mediation. There is no evidence showing EEG power could mediate sex effect on BOLD amplitude.

**Table 1.**
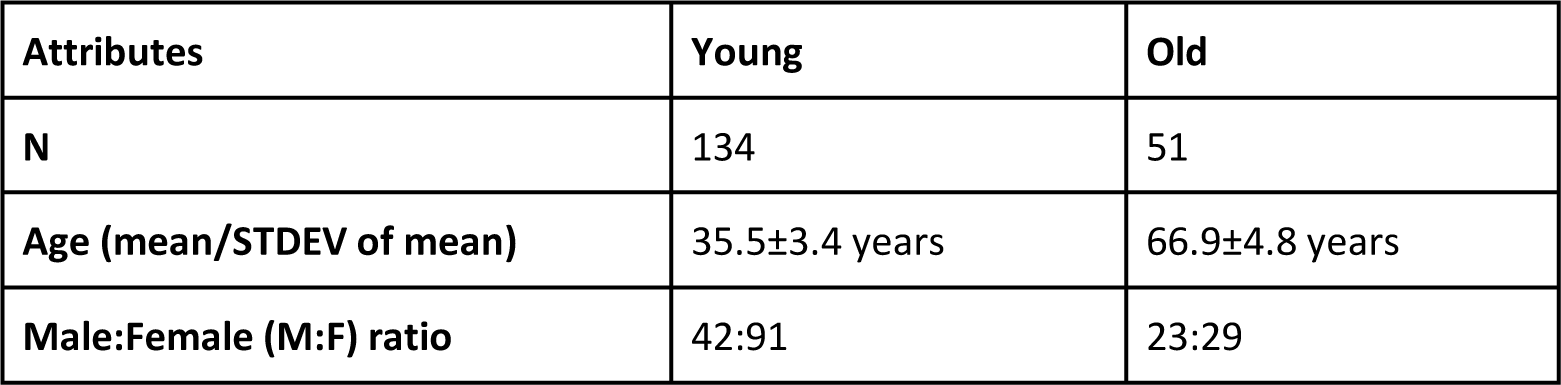
Subject demographics.

| Attributes | Young | Old |
| --- | --- | --- |
| **N** | 134 | 51 |
| **Age (mean/STDEV of mean)** | 35.5±3.4 years | 66.9±4.8 years |
| **Male:Female (M:F) ratio** | 42:91 | 23:29 |

#### Model 2: EEG frequency mediating the age or sex differences in fMRI frequency

There is no evidence showing EEG peak frequency could be the mediator for the BOLD peak frequency age and sex effect.

#### Model 3: EEG frequency mediating the age or sex differences in fMRI STDEV

There is no evidence showing EEG peak frequency could be the mediator for the age and sex associations in BOLD amplitude.

#### Comparison between epoch and non-epoch approaches

The use of the epoch-based pipeline was motivated by the desire to match the EEG and fMRI time scales. A comparison (using pairwise t-test) between the Sobel test (Sobel, M. E., 1982) derived p-value of epoch based and non-epoch based EEG parameters revealed that epoch-based alpha and beta band EEG power better mediate age effects on BOLD frequency, and delta EEG power better mediates age effects on BOLD amplitude in the 0.01-0.1 Hz band. For peak frequency, mediation percentages are lower in the epoch-based EEG signal between the age effect of BOLD low-frequency component peak frequency with the alpha band and beta band peak frequency, but higher between theta band frequency and BOLD frequency in the 0.1-0.3 Hz band. Over all, the mediation percentages are low and considered negligible.

## Discussion

Brain-signal variability, as measured using EEG, MEG or rs-fMRI, can all be indicative of functional integrity. It is widely assumed that fMRI signal fluctuations represent neuronal fluctuations, and there is still a limited amount of direct comparison of EEG and fMRI signal variability. In this work, we use the term “power” and “fluctuation amplitude” to refer to signal variability, with the intention of distinguishing it from age-related or sex-related variability. This study extends the previous study by Kumral et al. in several ways: (1) in addition to only investigating the standard deviation of fMRI and EEG signals, we also examine their characteristic frequencies; (2) we segregate the fMRI spectrum into a high- and a low-frequency band, each with a different neuronal weighting; (3) we examine age and sex effects on the ratio of fMRI and EEG signal power, a potential surrogate of vascular-neuronal coupling; (4) we use mediation analyses to investigate the associations between EEG and fMRI signal power and frequency. The main findings of this study are:

1. EEG power is generally higher in the young group except in the beta band.
2. EEG frequency is generally lower in the young group.
3. EEG power is higher in the female group across all bands, in many of the same regions that show age effects.
4. rs-fMRI signal fluctuation amplitudes are higher in young adults, most pronounced in the frequency > 0.1 Hz.
5. rs-fMRI signal fluctuation frequency is lower in young adults, also more pronounced in the frequency > 0.1 Hz.
6. rs-fMRI-EEG power ratio is higher in the young adults and lower in women.
7. Neither EEG power nor EEG frequency mediates the age and sex effects found in rs-fMRI fluctuation amplitude.

Thus, our first and third hypotheses are confirmed, but the second hypothesis is not.

### EEG versus age and sex

#### The effect of age on EEG power

The alpha band has been the most commonly studied, in part due to the ease of identifying the alpha peak. Slowing of alpha frequency has been associated with dementia (Garcés et al., 2013), but also with normal aging. Our findings of alpha power decline are consistent with the literature. Babiloni et al. (2006) demonstrated an age-dependent power decrease in alpha waves in the parietal, occipital and temporal regions (Babiloni et al., 2006). Magnetoencephalography (MEG) was used to report similar findings (Tsvetanov et al., 2015). On this topic, the review by Ishii et al., the details of which will not be repeated here, provides a broad summary of similar studies, demonstrating a largely posterior reduction in alpha power, which has in turn been associated with reductions in cognitive function (Ishii et al., 2017). Using the same LEMON data set, Kumral et al. reported a reduction in alpha power in the visual region. As expected, our results are in good agreement with Kumral et al., although our cohort was not selected in the same manner as in the Kumral study, which, in addition, used the finite element method for source reconstruction (in contrast to eLORETA).

Decreased delta rhythm in the occipital cortex has also been observed in aging using both EEG and MEG (Babiloni et al., 2006; Tsvetanov et al., 2015). We, like Kumral et al., found delta decline in aging to be found in the superior frontal gyrus, the precuneus, and the entire cingulate cortex. It should be noted that in the subcortical region, none but the delta band shows age effects in EEG power, and the effects cover most of the structures (including the thalamus, caudate, pallidum and hippocampus). In the theta band, as with the delta band, we observed age-associated power reduction in the superior frontal gyrus the precuneus, and the cingulate cortex, but with the addition of the superior parietal lobe. This is largely consistent with the findings of Kumral’s work, in which theta power was found to be lower in the older adults in the posterior default-mode network (Kumral et al., 2019). Frontal theta was spared of age effects, consistent with findings by Mizukami and Katada (Mizukami and Katada, 2018). Delta and theta power were previously found to positively associated with perception and executive function in older adults (Vlahou et al., 2014).

Our finding of age-related increase in beta power is consistent with the findings of Hashemi et al. in the frontal region (Hashemi et al., 2016), and are likely to be the result of the rise of beta1 and suppression of beta1 power in aging (Al Zoubi et al., 2018; Christov and Dushanova, 2016a). Furthermore, Kumral et al. also found an age-related enhancement in beta power in the temporal, pericentral and superior parietal regions, consistent with the spatial patterns identified in our findings.

### The effect of age on EEG frequency

In the epoch-based approach, the delta band shows higher frequency in young adults and theta band showing lower frequency in young adults. These observations are in part consistent with findings by Knyazeva et al., whereby low-frequency oscillations originating from the occipito-temporal regions in young adults move anteriorly with age (Knyazeva et al., 2018). In both epoch-based and non-epoch approaches, the beta band shows the strongest age effects (higher in older adults) in the cingulate and precuneus areas, which are generally associated with the default-mode network. The aging-related beta frequency elevation is consistent with the possible shift of sensory processing to high-frequency stimuli, leading to an increase in high-beta oscillations in older adults (Christov and Dushanova, 2016b). Moreover, regions showing beta frequency increases appear to not overlap at all with regions showing beta power increase in aging. Chiang et al. reported frontal-occipital alpha frequency differences (lower alpha frequency in frontal lobe) that may be altered by aging (Chiang et al., 2011), which, as Knyazeva et al. described, amounts to a merging of high- and low-frequency alpha peaks in aging (Knyazeva et al., 2018). However, this did not translate into a frequency shift for alpha. Lastly, there is no EEG frequency age effect in the subcortical region.

### The effect of sex on EEG

The literature is divided on the question of sex effects on resting EEG power (Zappasodi et al., 2006), but the majority of studies, especially those based on large data sets (Aurlien et al., 2004; Hashemi et al., 2016), show that EEG power is generally higher in women. In the study closest to ours, Kumral et al. found higher delta and theta power in the occipital-temporal cortex, higher alpha power in the frontal region and higher beta in the frontal as well as the occipital-temporal region in the female group (Kumral et al., 2019). All of these findings were reproduced by our study. Extending Kumral’s study, we found sex effects in all but the alpha band (female power > male), and in most of the subcortical regions. This finding is consistent with previous findings by Wada et al. (Wada et al., 1994), whereby the alpha band (high alpha only) also shows sex diffrences. Conversely, we found no strong association between sex and EEG frequency in any of the bands.

### Age-sex interaction

Kumral et al. reported that older adults show higher beta EEG power, driven by the female subjects. As we used a very similar cohort, we expected the same finding. However, we found that while men exhibit more age-group differences than women in the beta band, women exhibit more age differences in the theta and delta bands. Conversely, in the beta and delta bands, older adults exhibit greater sex differences than young adults. The differences with previous results could be attributed to differences in definitions of the EEG frequency bands. In Kumral et al.’s work, delta and beta bands are defined as 1-3 Hz and 15-25 Hz, respectively, whereas in our approach, delta is 1-4 Hz and beta is 12-30 Hz. The choice of a broader beta band is prompted by the large amount of beta power that still exists above 25 Hz, which may underlie the greater age difference in men, and will require further investigation in our future work.

### fMRI versus age and sex

#### The effect of age on fMRI amplitude

The BOLD signal fluctuation amplitude has been well studied in aging (Garrett et al., 2011, 2009; Guitart-Masip et al., 2016). Most studies of this type focus on the 0.01-0.1 Hz range, as it is most relevant to resting-state connectivity analysis. Like Grady and Garret (Grady and Garrett, 2014), we showed older adults have subdued inter-regional variability in their fMRI fluctuation amplitudes. The effect extends across the cortex as well as the putamen, similar to results illustrated by Grady and Garrett. Furthermore, reduced BOLD fluctuation amplitude has been associated with reduced cognitive performance and slower transition from rest to task states (Grady and Garrett, 2014). This is the finding that prompted us to try to associate fMRI age effects to those of EEG, as will be discussed in a later section. Unlike Mather et al. (Mather and Nga, 2013), we did not report a higher thalamic fluctuation amplitude in older adults, but we note Mather et al. examined fractional amplitude in the 0.01-0.1 Hz band, whereas we examined the quantitative EEG power.

In our study, in addition to the typically studied 0.01-0.1 Hz frequency band, we also studied the 0.1-0.3 Hz band. In this context, the most relevant previous study is by Yang et al. (Yang et al., 2018), whereby the rs-fMRI signal was categorized into high (0.087±0.2 Hz), low (0.045±0.087 Hz), and very-low (0.045 Hz) frequency bands using the Hilbert-Huang Transform (Yang et al., 2018). In the “low” frequency band, Yang et al. found reduced fMRI amplitude in the older adults. This is similar to our findings. As we found the fMRI signal in the 0.1-0.3 Hz frequency band to not produce the robust functional-connectivity patterns typically produced using the 0.01-0.1 Hz band (Yuen et al., 2019), we conclude, like Yang et al., that this high frequency band is dominated by non-neuronal effects such as respiration and head motion.

#### The effect of age on fMRI frequency

The significance of BOLD fluctuation frequency has been largely uninvestigated. To date, the most relevant study is by Yang et al. (Yang et al., 2018), who found aging-related IMF frequency increases in IMFs 1-5, which correspond to frequencies ranging from 0.045 to 0.1 Hz. This is consistent with our findings (for the 0.01-0.1 Hz band). However, we also found significant and widespread aging-related increases in frequency of the 0.1-0.3 Hz band, which is typically associated with an increased contribution from physiological processes. Interestingly, the most significant fMRI amplitude and frequency age effects, though in opposite directions, converge in the cingulate cortex.

Combined with the observation that overall fMRI power decreases with age in both frequency bands, we conclude that the apparent increase in frequency with age is due to a reduced contribution of low-frequency fluctuations. In the 0.01-0.1 Hz band, the low-frequency contribution can result from the contributions of arterial carbon dioxide (CO2) and heart-rate variability (HRV) to the signal. Very-low frequency vascular oscillations measured using near-infrared spectroscopy have been found to decline with age (Schroeter et al., 2004), and coincide in frequency with the resting vascular response to arterial carbon-dioxide fluctuations (0.02-0.04 Hz) (Golestani et al., 2015; Liu et al., 2017). In addition, low-frequency HRV, spanning 0.04-0.15 Hz, is found to decrease in aging (Rizzo et al., 1999), particularly in women (Moodithaya and Avadhany, 2012). In the 0.1-0.3 Hz band, the lower-frequency contribution is likely to stem from the neuronally-driven fluctuations centred at 0.1 Hz. This is the frequency band typically used for functional-connectivity mapping, and a reduction in fMRI signal amplitude in this range would be consistent with similar findings in the EEG data. Also in this frequency band are vasomotion and Mayer waves, which are found to decline in aging as well, consistent with reduced vascular smooth-muscle activity and increased vascular stiffening (Schroeter et al., 2004). Moreover, it is likely that older adults may express higher head motion (van Dijk et al., 2011), adding to the age-related shift of the fMRI signal towards higher frequency content. Like Kumral et al, we found no significant effect of sex on fMRI amplitude (Kumral et al., 2019). We also failed to detect an effect on fMRI frequency.

The effect of sex on fMRI amplitude and frequency measures is negligible, in contrast to findings in EEG.

### Resting-state fMRI-EEG associations

Younger adults are found to have higher fMRI-EEG power ratios. The effects are most pronounced in the beta band, and the affected regions span nearly the entire anterior cortex, also including subcortical regions such as the putamen and pallidum. Moreover, the age effect on the ratio is more pronounced in the 0.1-0.3 Hz range than in the 0.01-0.1 Hz range. This can be potentially be explained by two aspects: (1) the fMRI-EEG ratio in the 0.01-0.1 Hz band reflects neurovascular coupling ratio, known to be lower in older adults (Tarantini et al., 2017); (2) the fMRI-EEG ratio in the 0.1-0.3 Hz band reflects fluctuations in the fMRI signal with minimal corresponding fluctuations in the EEG signal, suggestive of non-neuronal origins of such fMRI fluctuations. Nonetheless, the age effects on the fMRI-EEG power ratios overlap highly between the 0.01-0.1 Hz and 0.1-0.3 Hz bands, suggesting related mechanisms for these age effects. One possible source of this is that age-related Fmri reductions at 0.02-0.04 Hz (due to vascular reactivity decline) and reductions at 0.1 Hz (due to neural-activity decline) coexist.

Significant sex effects (female < male) are found in the ratios corresponding to all EEG bands, spanning the frontal, occipital and paracentral regions. The effects are greatest for the beta band, and more widespread for the 0.1-0.3 Hz fMRI-beta ratio. This is to be expected, as beta power is significantly higher in women, so long as the fMRI signal amplitude remains unaffected by beta power (as is the case). These results also suggest that the sex differences in the 0.1-0.3 Hz band is greater than in the 0.01-0.1 Hz band, although in the study these differences were not found to be significant.

Given the general consensus that fMRI signal and evoked EEG potentials can be associated through a hemodynamic response function (de Munck et al., 2007; Goldman et al., 2002), one might assume that resting-state fMRI and EEG power as well as frequency to be associated. Portnova et al. reported that resting delta power is strongly correlated with rs-fMRI signal amplitude in the bilateral parahippocampal gyri, middle frontal gyri (Portnova et al., 2017). Specific to the effect of aging, although both EEG and fMRI power are both lower in the older adults (with the exception of the beta band), and both display peak age effects in the cingulate and precuneus, the mediation effect of EEG (amplitude or frequency) on fMRI fluctuation amplitude was negligible. Consistent with this finding, Kumral et al. found no correlation between EEG power and fMRI amplitude in the 0.01-0.1Hz band (Kumral et al., 2019). The alternative mediator, as indicated by the study of Tsvetanov et al., would be vascular reactivity (Tsvetanov et al., 2015). Based on observations in mild-cognitive impairment, aging is likely to be associated with a lower and slower vascular response (Richiardi et al., 2015). Moreover, arterial dysfunction is observed in early aging in the mouse model (Balbi et al., 2015). Nonetheless, Garrett et al. demonstrated that rs-fMRI amplitude effects in aging are not solely due to vascular differences between age groups (Garrett et al., 2017). Thus, we conclude that rs-fMRI amplitude is a potential imaging marker that contains unique effects of aging.

### Limitations

As the LEMON EEG data does not contain gamma band EEG, we limited our analysis to alpha, beta, delta and theta bands. Moreover, the EEG and fMRI data were not acquired simultaneously. While this ensured superior data quality of both modalities, it precluded us from some interesting dynamic analyses. Finally, the sex ratios are different between young and old groups. Nonetheless, we performed sex-specific age analysis (and age-specific sex analysis) to examine sex-related biases.

## Conclusions

To conclude, EEG power and frequency are both lower in the older adults except in the beta band, where both are higher in the older group. There was no spatial association between power and frequency in each EEG band, and none of the EEG metrics mediated the age or sex effects on fMRI fluctuation amplitude. Finally, the fMRI-EEG power ratio can be an interesting marker of the physiological process involved in brain aging, especially given its selective age sensitivity to the beta EEG band.

## Acknowledgments

The authors acknowledge funding support from the Canadian Institutes of Health Research (CIHR) and the Natural Sciences and Engineering Research Council of Canada (NSERC). We are also grateful to Mr. Jacob Matthews, Mr. Nick Teller and Mr. Jordan Chad for proofreading.

## Notes

### Competing Interest Statement

The authors have declared no competing interest.

## References

Al Zoubi, O., Ki Wong, C., Kuplicki, R.T., Yeh, H.-W., Mayeli, A., Refai, H., Paulus, M., Bodurka, J., 2018. Predicting Age From Brain EEG Signals-A Machine Learning Approach. Front. Aging Neurosci. 10, 184.

Aurlien, H., Gjerde, I.O., Aarseth, J.H., Eldøen, G., Karlsen, B., Skeidsvoll, H., Gilhus, N.E., 2004. EEG background activity described by a large computerized database. Clin. Neurophysiol. 115, 665–673.

Babayan, A., Erbey, M., Kumral, D., Reinelt, J.D., Reiter, A.M.F., Röbbig, J., Schaare, H.L., Uhlig, M., Anwander, A., Bazin, P.-L., Horstmann, A., Lampe, L., Nikulin, V.V., Okon-Singer, H., Preusser, S., Pampel, A., Rohr, C.S., Sacher, J., Thöne-Otto, A., Trapp, S., Nierhaus, T., Altmann, D., Arelin, K., Blöchl, M., Bongartz, E., Breig, P., Cesnaite, E., Chen, S., Cozatl, R., Czerwonatis, S., Dambrauskaite, G., Dreyer, M., Enders, J., Engelhardt, M., Fischer, M.M., Forschack, N., Golchert, J., Golz, L., Guran, C.A., Hedrich, S., Hentschel, N., Hoffmann, D.I., Huntenburg, J.M., Jost, R., Kosatschek, A., Kunzendorf, S., Lammers, H., Lauckner, M.E., Mahjoory, K., Kanaan, A.S., Mendes, N., Menger, R., Morino, E., Näthe, K., Neubauer, J., Noyan, H., Oligschläger, S., Panczyszyn-Trzewik, P., Poehlchen, D., Putzke, N., Roski, S., Schaller, M.-C., Schieferbein, A., Schlaak, B., Schmidt, R., Gorgolewski, K.J., Schmidt, H.M., Schrimpf, A., Stasch, S., Voss, M., Wiedemann, A., Margulies, D.S., Gaebler, M., Villringer, A., 2019. A mind-brain-body dataset of MRI, EEG, cognition, emotion, and peripheral physiology in young and old adults. Sci Data 6, 180308.

Babiloni, C., Binetti, G., Cassarino, A., Forno, G.D., Del Percio, C., Ferreri, F., Ferri, R., Frisoni, G., Galderisi, S., Hirata, K., Lanuzza, B., Miniussi, C., Mucci, A., Nobili, F., Rodriguez, G., Romani, G.L., Rossini, P.M., 2006. Sources of cortical rhythms in adults during physiological aging: A multicentric EEG study. Human Brain Mapping. https://doi.org/10.1002/hbm.20175

Balbi, M., Ghosh, M., Longden, T.A., Jativa Vega, M., Gesierich, B., Hellal, F., Lourbopoulos, A., Nelson, M.T., Plesnila, N., 2015. Dysfunction of mouse cerebral arteries during early aging. J. Cereb. Blood Flow Metab. 35, 1445–1453.

Baron, R.M., Kenny, D.A., 1986. The moderator–mediator variable distinction in social psychological research: Conceptual, strategic, and statistical considerations. J. Pers. Soc. Psychol. 51, 1173–1182.

Bell, A.J., Sejnowski, T.J., 1997. The “independent components” of natural scenes are edge filters. Vision Res. 37, 3327–3338.

Chiang, A.K.I., Rennie, C.J., Robinson, P.A., van Albada, S.J., Kerr, C.C., 2011. Age trends and sex differences of alpha rhythms including split alpha peaks. Clinical Neurophysiology. https://doi.org/10.1016/j.clinph.2011.01.040

Christov, M., Dushanova, J., 2016a. Functional correlates of brain aging: beta and gamma components of event-related band responses. Acta Neurobiologiae Experimentalis. https://doi.org/10.21307/ane-2017-009

Christov, M., Dushanova, J., 2016b. Functional Correlates of the Aging Brain: Beta Frequency Band Responses to Age-related Cortical Changes. International Journal of Neurorehabilitation. https://doi.org/10.4172/2376-0281.1000194

Daunizeau, J., Adam, V., Rigoux, L., 2014. VBA: a probabilistic treatment of nonlinear models for neurobiological and behavioural data. PLoS Comput. Biol. 10, e1003441.

de Munck, J.C., Gonçalves, S.I., Huijboom, L., Kuijer, J.P.A., Pouwels, P.J.W., Heethaar, R.M., Lopes da Silva, F.H., 2007. The hemodynamic response of the alpha rhythm: an EEG/fMRI study. Neuroimage 35, 1142–1151.

Desikan, R.S., Ségonne, F., Fischl, B., Quinn, B.T., Dickerson, B.C., Blacker, D., Buckner, R.L., Dale, A.M., Maguire, R.P., Hyman, B.T., Albert, M.S., Killiany, R.J., 2006. An automated labeling system for subdividing the human cerebral cortex on MRI scans into gyral based regions of interest. Neuroimage 31, 968–980.

Garcés, P., Vicente, R., Wibral, M., Pineda Pardo, J.A., Lopez, M.E., Aurtenetxe, S., 2013. Brain-wide slowing of spontaneous alpha rhythms in mild cognitive impairment. Front. Aging Neurosci. 7, 100.

Garrett, D.D., Kovacevic, N., McIntosh, A.R., Grady, C.L., 2010. Blood oxygen level-dependent signal variability is more than just noise. J. Neurosci. 30, 4914–4921.

Garrett, D.D., Kovacevic, N., McIntosh, A.R., Grady, C.L., 2009. The Importance of Brain Variability. NeuroImage. https://doi.org/10.1016/s1053-8119(09)70579-8

Garrett, D.D., Lindenberger, U., Hoge, R.D., Gauthier, C.J., 2017. Age differences in brain signal variability are robust to multiple vascular controls. Sci. Rep. 7, 10149.

Garrett, D.D., McIntosh, A.R., Grady, C.L., 2011. Moment-to-moment signal variability in the human brain can inform models of stochastic facilitation now. Nat. Rev. Neurosci.

Goldman, R.I., Stern, J.M., Engel, J., Cohen, M.S., 2002. Simultaneous EEG and fMRI of the alpha rhythm. NeuroReport. https://doi.org/10.1097/00001756-200212200-00022

Golestani, A.M., Chang, C., Kwinta, J.B., Khatamian, Y.B., Chen, J.J., 2015. Mapping the end-tidal CO2 response function in the resting-state BOLD fMRI signal: Spatial specificity, test–retest reliability and effect of fMRI sampling rate. Neuroimage 104, 266–277.

Grady, C.L., Garrett, D.D., 2014. Understanding variability in the BOLD signal and why it matters for aging. Brain Imaging Behav. 8, 274–283.

Guitart-Masip, M., Salami, A., Garrett, D., Rieckmann, A., Lindenberger, U., Bäckman, L., 2016. BOLD Variability is Related to Dopaminergic Neurotransmission and Cognitive Aging. Cereb. Cortex 26, 2074–2083.

Hashemi, A., Pino, L.J., Moffat, G., Mathewson, K.J., Aimone, C., Bennett, P.J., Schmidt, L.A., Sekuler, A.B., 2016. Characterizing Population EEG Dynamics throughout Adulthood. eneuro. https://doi.org/10.1523/eneuro.0275-16.2016

Haufe, S., Ewald, A., 2019. A Simulation Framework for Benchmarking EEG-Based Brain Connectivity Estimation Methodologies. Brain Topogr. 32, 625–642.

Hayes, A.F., 2013. Introduction to mediation, moderation, and conditional process analysis: A regression-based approach. Methodology in the social sciences. 507.

Hjelmervik, H., Hausmann, M., Osnes, B., Westerhausen, R., Specht, K., 2014. Resting States Are Resting Traits – An fMRI Study of Sex Differences and Menstrual Cycle Effects in Resting State Cognitive Control Networks. PLoS ONE. https://doi.org/10.1371/journal.pone.0103492

Ishii, R., Canuet, L., Aoki, Y., Hata, M., Iwase, M., Ikeda, S., Nishida, K., Ikeda, M., 2017. Healthy and Pathological Brain Aging: From the Perspective of Oscillations, Functional Connectivity, and Signal Complexity. Neuropsychobiology 75, 151–161.

Jiao, F., Gao, Z., Shi, K., Jia, X., Wu, P., Jiang, C., Ge, J., Su, H., Guan, Y., Shi, S., Zang, Y.-F., Zuo, C., 2019. Frequency-Dependent Relationship Between Resting-State fMRI and Glucose Metabolism in the Elderly. Front. Neurol. 10, 566.

Kannurpatti, S.S., Rypma, B., Biswal, B.B., 2012. Prediction of Task-Related BOLD fMRI with Amplitude Signatures of Resting-State fMRI. Front. Syst. Neurosci. 6, 7.

Knyazeva, M.G., Barzegaran, E., Vildavski, V.Y., Demonet, J.-F., 2018. Aging of human alpha rhythm. Neurobiol. Aging 69, 261–273.

Kumral, D., Şansal, F., Cesnaite, E., Mahjoory, K., Al, E., Gaebler, M., Nikulin, V.V., Villringer, A., 2019. BOLD and EEG signal variability at rest differently relate to aging in the human brain. Neuroimage 116373.

Liu, P., Li, Y., Pinho, M., Park, D.C., Welch, B.G., Lu, H., 2017. Cerebrovascular reactivity mapping without gas challenges. Neuroimage 146, 320–326.

Mather, M., Nga, L., 2013. Age differences in thalamic low-frequency fluctuations. NeuroReport. https://doi.org/10.1097/wnr.0b013e32835f6784

Mizukami, K., Katada, A., 2018. EEG Frequency Characteristics in Healthy Advanced Elderly. Journal of Psychophysiology. https://doi.org/10.1027/0269-8803/a000190

Moodithaya, S., Avadhany, S.T., 2012. Gender differences in age-related changes in cardiac autonomic nervous function. J. Aging Res. 2012, 679345.

Pascual-Marqui, R.D., 2007. Discrete, 3D distributed, linear imaging methods of electric neuronal activity. Part 1: exact, zero error localization. arXiv [math-ph].

Portnova, G.V., Tetereva, A., Balaev, V., Atanov, M., Skiteva, L., Ushakov, V., Ivanitsky, A., Martynova, O., 2017. Correlation of BOLD Signal with Linear and Nonlinear Patterns of EEG in Resting State EEG-Informed fMRI. Front. Hum. Neurosci. 11, 654.

Richiardi, J., Monsch, A.U., Haas, T., Barkhof, F., Van de Ville, D., Radü, E.W., Kressig, R.W., Haller, S., 2015. Altered cerebrovascular reactivity velocity in mild cognitive impairment and Alzheimer’s disease. Neurobiol. Aging 36, 33–41.

Rizzo, V., Villatico Campbell, S., Di Maio, F., Tallarico, D., Lorido, A., Petretto, F., Bianchi, A., Carmenini, G., 1999. Spectral analysis of heart rate variability in elderly non-dipper hypertensive patients. Journal of Human Hypertension. https://doi.org/10.1038/sj.jhh.1000810

Rosenblum, M., Pikovsky, A., Kurths, J., Schäfer, C., Tass, P.A., 2001. Chapter 9 Phase synchronization: From theory to data analysis, in: Moss, F., Gielen, S. (Eds.), Handbook of Biological Physics. North-Holland, pp. 279–321.

Schroeter, M.L., Schmiedel, O., von Cramon, D.Y., 2004. Spontaneous Low-Frequency Oscillations Decline in the Aging Brain. Journal of Cerebral Blood Flow & Metabolism. https://doi.org/10.1097/01.wcb.0000135231.90164.40

Sobel, M.E., 1982. Asymptotic Confidence Intervals for Indirect Effects in Structural Equation Models. Sociol. Methodol. 13, 290–312.

Tarantini, S., Tran, C.H.T., Gordon, G.R., Ungvari, Z., Csiszar, A., 2017. Impaired neurovascular coupling in aging and Alzheimer’s disease: Contribution of astrocyte dysfunction and endothelial impairment to cognitive decline. Exp. Gerontol. 94, 52–58.

Tsvetanov, K.A., Henson, R.N.A., Tyler, L.K., Davis, S.W., Shafto, M.A., Taylor, J.R., Williams, N., Cam-Can Rowe, J.B., 2015. The effect of ageing on fMRI: Correction for the confounding effects of vascular reactivity evaluated by joint fMRI and MEG in 335 adults. Hum. Brain Mapp. 36, 2248–2269.

van Dijk, K.R.A., Sabuncu, M.R., Buckner, R.L., 2011. The influence of head motion on intrinsic functional connectivity MRI. Neuroimage 59, 431–438.

Vlahou, E.L., Thurm, F., Kolassa, I.-T., Schlee, W., 2014. Resting-state slow wave power, healthy aging and cognitive performance. Sci. Rep. 4, 5101.

Wada, Y., Takizawa, Y., Zheng-Yan, J., Yamaguchi, N., 1994. Gender Differences in Quantitative EEG at Rest and during Photic Stimulation in Normal Young Adults. Clinical Electroencephalography. https://doi.org/10.1177/155005949402500209

Yang, A.C., Tsai, S.-J., Lin, C.-P., Peng, C.-K., Huang, N.E., 2018. Frequency and amplitude modulation of resting-state fMRI signals and their functional relevance in normal aging. Neurobiol. Aging 70, 59–69.

Yuen, N.H., Osachoff, N., Chen, J.J., 2019. Intrinsic Frequencies of the Resting-State fMRI Signal: The Frequency Dependence of Functional Connectivity and the Effect of Mode Mixing. Front. Neurosci. 13, 900.

Zappasodi, F., Pasqualetti, P., Tombini, M., Ercolani, M., Pizzella, V., Rossini, P.M., Tecchio, F., 2006. Hand cortical representation at rest and during activation: Gender and age effects in the two hemispheres. Clinical Neurophysiology. https://doi.org/10.1016/j.clinph.2006.03.016

Zhang, C., Cahill, N.D., Arbabshirani, M.R., White, T., Baum, S.A., Michael, A.M., 2016. Sex and Age Effects of Functional Connectivity in Early Adulthood. Brain Connectivity. https://doi.org/10.1089/brain.2016.0429

Zou, Q.H., Zhu, C.Z., Yang, Y., Zuo, X.N., Long, X.Y., Cao, Q.J., Wang, Y.F., Zang, Y.F., 2008. An improved approach to detection of amplitude of low-frequency fluctuation (ALFF) for resting-state fMRI: fractional ALFF. J. Neurosci. Methods 172, 137–141.

